# Sequencing of Argonaute-bound miRNA/mRNA hybrids reveals regulation of the unfolded protein response by microRNA-320a

**DOI:** 10.1101/2021.10.05.463240

**Authors:** Christopher J. Fields, Lu Li, Nicholas M. Hiers, Tianqi Li, Peike Sheng, Taha Huda, Jixiu Shan, Lauren Gay, Tongjun Gu, Jiang Bian, Michael S. Kilberg, Rolf Renne, Mingyi Xie

## Abstract

MicroRNAs (miRNA) are short non-coding RNAs widely implicated in gene regulation. Most metazoan miRNAs utilize the RNase III enzymes Drosha and Dicer for biogenesis. One notable exception is the RNA polymerase II transcription start sites (TSS) miRNAs whose biogenesis does not require Drosha. The functional importance of the TSS-miRNA biogenesis is uncertain. To better understand the function of TSS-miRNAs, we applied a modified <u>C</u>rosslinking, <u>L</u>igation, and <u>S</u>equencing of <u>H</u>ybrids on Argonaute (AGO-qCLASH) to identify the targets for TSS-miRNAs in HCT116 colorectal cancer cells with or without *DROSHA* knockout. We observed that miR-320a hybrids dominate in TSS-miRNA hybrids identified by AGO-qCLASH. Targets for miR-320a are enriched in the eIF2 signaling pathway, a downstream component of the unfolded protein response. Consistently, in miR-320a mimic- and antagomir- transfected cells, differentially expressed genes are enriched in eIF2 signaling. Within the AGO-qCLASH data, we identified the endoplasmic reticulum (ER) chaperone Calnexin as a direct miR-320a target, thus connecting miR-320a to the unfolded protein response. During ER stress, but not amino acid deprivation, miR-320a up-regulates ATF4, a critical transcription factor for resolving ER stress. Our study investigates the targetome of the TSS-miRNAs in colorectal cancer cells and establishes miR-320a as a regulator of unfolded protein response.

## INTRODUCTION

MicroRNAs (miRNAs) are ~18-22 nucleotide non-coding RNAs that target the majority of messenger RNAs (mRNAs) for post-transcriptional regulation in a tissue-specific manner (1). Biogenesis of miRNAs begins in the nucleus, where primary miRNA (pri-miRNA) is transcribed and folds into a stem-loop hairpin that is subsequently recognized by the Microprocessor (2, 3). The Microprocessor complex comprises two RNA-binding proteins (DGCR8) and one RNase III enzyme (Drosha) (4). Following Drosha cleavage, Exportin-5 shuttles the resulting precursor miRNA (pre-miRNA) into the cytoplasm, where Dicer removes the loop to form a ~22 base pair double-stranded RNA (5). Mature miRNA is loaded onto the effector protein Argonaute (AGO), forming the RNA-induced silencing complex (RISC) (5, 6). One strand is retained as part of the RISC and guides the complex to target mRNAs. Canonically, miRNAs target mRNA’s 3′ untranslated region (UTR) using their seed sequence (nucleotides 2-8) by Watson-Crick base-pairing (7). The binding of the RISC complex leads to the recruitment of the CCR4-NOT deadenylase and DCP1-DCP2 decapping complexes for mRNA degradation (8).

While the mechanisms of canonical miRNA biogenesis are well described, the existence and importance of Dicer- or Drosha-independent modes of processing is increasingly appreciated (9, 10). One such pathway is the Dicer-independent processing of miR-451, in which AGO2 directly slices the precursor hairpin of miR-451 following its export from the cytoplasm (11–13). While the majority of the miRNAs are increasingly down-regulated during erythrocyte differentiation, the Dicer-independent miR-451 is up-regulated (14, 15). Likewise, two distinct pathways that bypass Drosha have been identified. The mirtron pathway generates pre-miRNAs from intron splicing and then converges into the canonical pathway using nuclear-cytoplasmic transport by Exportin-5 (16, 17). The transcription start sites (TSS)-miRNA pathway directly generates pre-miRNAs at RNA polymerase II transcription start sites, negating the need for Drosha processing (18–20). Consequently, this processing results in the pre-miRNA with a 7-methylguanosine (m^7^G) cap on the 5′ end instead of the canonical 5′ monophosphate (18). Moreover, some TSS-miRNA precursors contain extensions in their 5′ ends (20). These TSS-miRNAs could have unique functions in development and disease, especially where the Microprocessor activity is perturbed (21).

Previously, we observed that TSS-miRNAs are enriched in HCT116 *DROSHA* knockout (KO) cells compared to wild-type (WT) cells, likely due to better access to the limited miRNA machinery (Fig. 1A) (20, 22). Because TSS-miRNAs are synthesized independently of the Microprocessor, it is conceivable that they persist in diseased cell types where *DROSHA* function is impaired. The loss of *DROSHA* has been documented in both endometrial cancer and breast cancer (23, 24). In melanoma, Drosha levels in the nucleus are reduced, likely due to impaired cytoplasmic-nuclear import (25). Multiple components of the miRNA biogenesis pathway are mutated in pediatric Wilms tumors, including *DROSHA* which is down-regulated in around 12% of cases (26–28). Accordingly, in the Wilms tumor cases with lower *DROSHA* expression, TSS-miRNAs, including miR-320a, miR-484, and mirtron miR-877 are up-regulated (26, 29). The consequences of *DROSHA* down-regulation and the corresponding up-regulation of TSS-miRNAs are poorly understood due to the complexity and incompleteness of the TSS-miRNA targetome.

**Figure 1.**
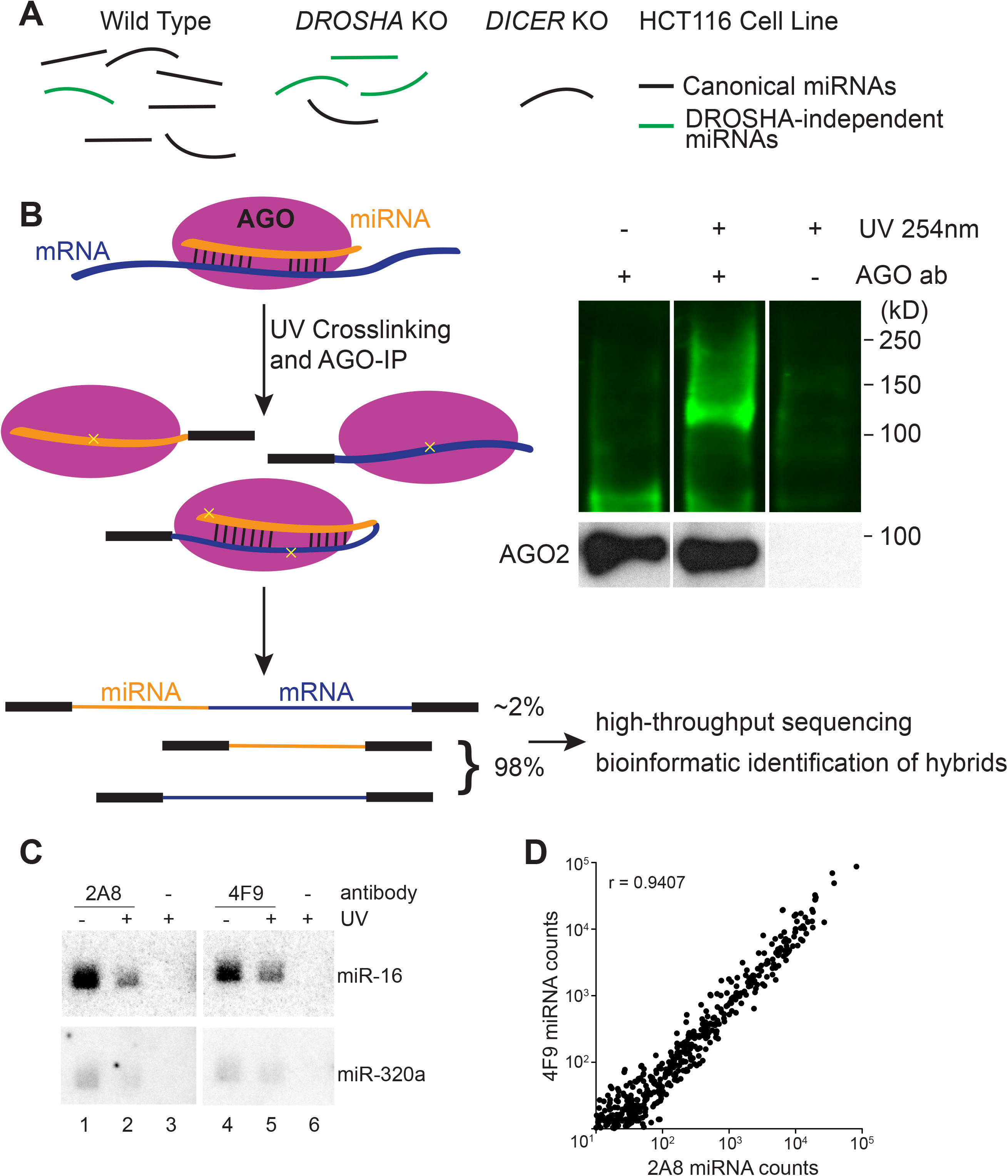
Argonaute crosslinking, ligation, and sequencing of hybrids in colorectal cancer cells. (A) HCT116 cell lines containing knockouts for core miRNA processing machinery allow the enrichment of Drosha-independent miRNAs (e.g., mirtrons, TSS-miRNAs). (B) Schematic of AGO-CLASH. In both CLASH and qCLASH, endogenous miRNA and mRNA are crosslinked (yellow crosses) and immunoprecipitated using AGO antibody. Following AGO-IP, the miRNA is ligated directly to its targets to form a single hybrid molecule. Each library produces hybrid reads in addition to individual reads. Blots show AGO-RNA complexes detected on nitrocellulose membrane using IRDye 800CW. AGO-bound ribonucleic molecules were ligated with RA3 primer containing an IRDye, separated on a NuPAGE Bis-Tris gel, and transferred to nitrocellulose. (C) ^32^P Northern blot of AGO-bound RNAs in HCT116 cells probed for miR-16 and miR-320a. (D) Non-hybrid miRNA reads from qCLASH libraries were quantitated with miR-Deep2. Normalized counts from qCLASH libraries that used the same AGO antibody were averaged, showing that both 2A8 and 4F9 pull down similar amounts of each miRNA. The correlation coefficient (r) is depicted on the graph.

Transcriptome-wide miRNA binding sites can be experimentally identified by crosslinking and immunoprecipitation of AGO in tandem with high throughput sequencing (e.g., HiTS-CLIP, PAR-CLIP) (30, 31). Compared with pure bioinformatic prediction algorithms, AGO-CLIP offers the advantage of defining the miRNA targetome in cell-specific contexts. Still, AGO-CLIP protocols are laborious and require bioinformatic prediction of miRNA/target pairing, resulting in a large percentage of target sites that do not confer effective repression when bound by miRNAs (32). Others have improved upon AGO-CLIP by directly ligating miRNAs to targets while they are base-pairing inside AGO. Such crosslinking, ligation, and sequencing of hybrids (AGO-CLASH, referred to as CLASH hereafter for simplicity) physically connects miRNA and target mRNA, allowing for high-confidence identification of the miRNA targetome (33, 34). Further iterations of CLASH, termed quick CLASH (qCLASH), have taken advantage of recent advances in bioinformatic approaches and the unprecedented depth offered by high-throughput sequencing technology. By omitting SDS-PAGE separation and purification of the protein-bound RNAs from nitrocellulose membrane, the qCLASH protocol is easier to adapt for both research and clinical purposes 35, 36).

Here, we identify the targets for TSS-miRNAs using qCLASH in colorectal cancer HCT116 cells with or without Drosha. We discovered miR-320a to be the most abundant TSS-miRNA in HCT116 cells and verified that it represses targets identified by qCLASH. Pathway analysis revealed that miR-320a affects the integrated stress response (ISR), one of three pathways that collectively make up the endoplasmic reticulum (ER) unfolded protein response (UPR). Specifically, qCLASH identified the ER chaperone Calnexin (CANX) as a top miR-320a target in WT and *DROSHA* KO cells. Both *CANX* mRNA and protein abundance were repressed by miR-320a. The targeting of *CANX* by miR-320a activates the ISR, a process that plays an essential role in maintaining homeostasis in the ER. Furthermore, we demonstrated that miR-320a directly targets the qCLASH-identified binding sites in the 3′ UTR of *CANX*. Consistent with previous reports, miR-320a is repressed in colorectal cancer while *CANX* is up-regulated. Investigating the downstream consequences of miR-320a perturbation of the ISR, we discovered that miR-320a can enhance *ATF4* activation in cells during ER stress. In summary, we used qCLASH to identify the targetome of miR-320a in colorectal cancer cells and established that miR-320a regulates the ISR by targeting *CANX*.

## RESULTS

### CLASH for miRNA target identification in HCT116 cells

We used CLASH and its variant qCLASH to elucidate the miRNA targetomes in HCT116 cells, including WT, *DROSHA* KO, and *DICER* KO cells (Figs. 1A and 1B). *DROSHA* KO cells have elevated expression of Drosha-independent miRNAs, including TSS-miRNAs (20, 37). miRNA expression in *DICER* KO cells is abated and can serve as a background control. To minimize potential antibody specific bias, we used two different pan-AGO antibodies that can immunoprecipitate all four human AGO proteins (AGO1-AGO4) for CLASH and qCLASH. AGO antibody clone 2A8 has been widely used in the field for AGO-CLIP experiments, while clone 4F9 was less well characterized, but is locally available through the Interdisciplinary Center for Biotechnology Research at the University of Florida (30, 38–41). We found that clone 4F9 could immunoprecipitate AGO2 as well as the clone 2A8 (Fig. S1A, lanes 3 and 7). Next, we examined AGO-associated miRNAs that are enriched by the two different antibodies. Northern blot analyses showed that canonical miRNA (miR-16) and TSS-miRNA (miR-320a) could be efficiently isolated from cells with or without 254 nm ultraviolet (UV light) crosslinking using either antibody (Fig. 1C). Furthermore, high-throughput sequencing experiments revealed that miRNAs co-precipitated with AGO by both antibodies are highly correlated (r = 0.9407; Fig. 1D). Therefore, we concluded that the efficacy of the 4F9 antibody rivals the widely used 2A8 antibody, and thus both are suitable for CLASH experiments.

Following immunoprecipitation, AGO-bound RNAs were digested with RNase A, modified at the termini by T4 polynucleotide kinase, and finally intermolecularly ligated together with RNA ligase, forming a single hybrid molecule (see Materials and Methods for details). For both the CLASH and qCLASH protocols, we ligated a custom pre-adenylated 3′ adapter (Table S1). Compared to qCLASH, the CLASH protocol includes SDS-PAGE to separate crosslinked AGO-RNA complexes, which are then transferred to a nitrocellulose membrane and excised according to the appropriate molecular weight (Figs. 1B and S1B). For both CLASH and qCLASH protocols, RNAs were purified from protein complexes, ligated with 5′ adapter, reverse transcribed, and PCR amplified to be compatible with Illumina sequencing (Fig. S1C).

Paired-end sequencing reads obtained from qCLASH were filtered using Trimmomatic to remove low-quality reads and adapter sequences (42). Forward and reverse reads were combined using Pear to form a single sequence for analysis (43). Because the 5′ and 3′ adapters contain degenerate nucleotides to account for PCR bias and minimize ligation preferences (44), we collapsed duplicate reads using the FASTX-Toolkit FASTQ Collapser, and the degenerate nucleotides were removed using Cutadapt (45, 46). Chimeric reads (hybrids) were identified using the Hyb software with the default database supplemented with previously identified TSS-miRNAs and miR-snaR (20, 47, 48). With viennad-format files output from Hyb, we used a custom script to identify overlapping targets for specific miRNAs. CLASH libraries were analyzed as described above, except that FASTX-Toolkit FASTQ Collapser and Cutadapt were not used since the adapters used for these libraries do not contain random nucleotides.

### Comparison of CLASH, qCLASH, and miRNA-specific qCLASH

We initially performed a single biological replicate of CLASH and qCLASH using 2A8 AGO antibody in each HCT116 cell line (WT, *DROSHA* KO, and *DICER* KO). For CLASH, two regions that correspond to a molecular weight of 110-125 kD and 125-150 kD, which contain AGO crosslinked with different lengths of RNAs, were excised from the membrane (Fig. S1B). As expected, in data sets obtained from *DROSHA* KO cells, we observed enrichment of hybrids containing Drosha-independent miRNAs including TSS-miRNAs (miR-320a and miR-484), mirtron (miR-877), and the recently described miR-snaR, a non-canonical miRNA derived from non-coding RNA snaR-A (Table S2) (48). MiR-320a-containing hybrids are far more abundant than other Drosha-independent miRNAs, making up to 74.58% of all the miRNA/mRNA hybrids in qCLASH data (Table S2). Overall, with comparable amounts of raw reads obtained from both WT and *DROSHA* KO cells, we identified approximately four times more miRNA/mRNA hybrids in qCLASH compared with CLASH (70,124 : 16,487 for WT, and 4,752 : 1558 for *DROSHA* KO) (Table S2). For miR-320a specifically, qCLASH hybrids are six to eight times more than CLASH hybrids (4,511 : 543 for WT, and 3544 : 597 for *DROSHA* KO) (Table S2). Notably, approximately 50% of the CLASH hybrids can be found in qCLASH hybrids (Figs. 2A and 2B). It appears that more miRNA targeting interactions could be detected by qCLASH compared to CLASH.

**Figure 2:**
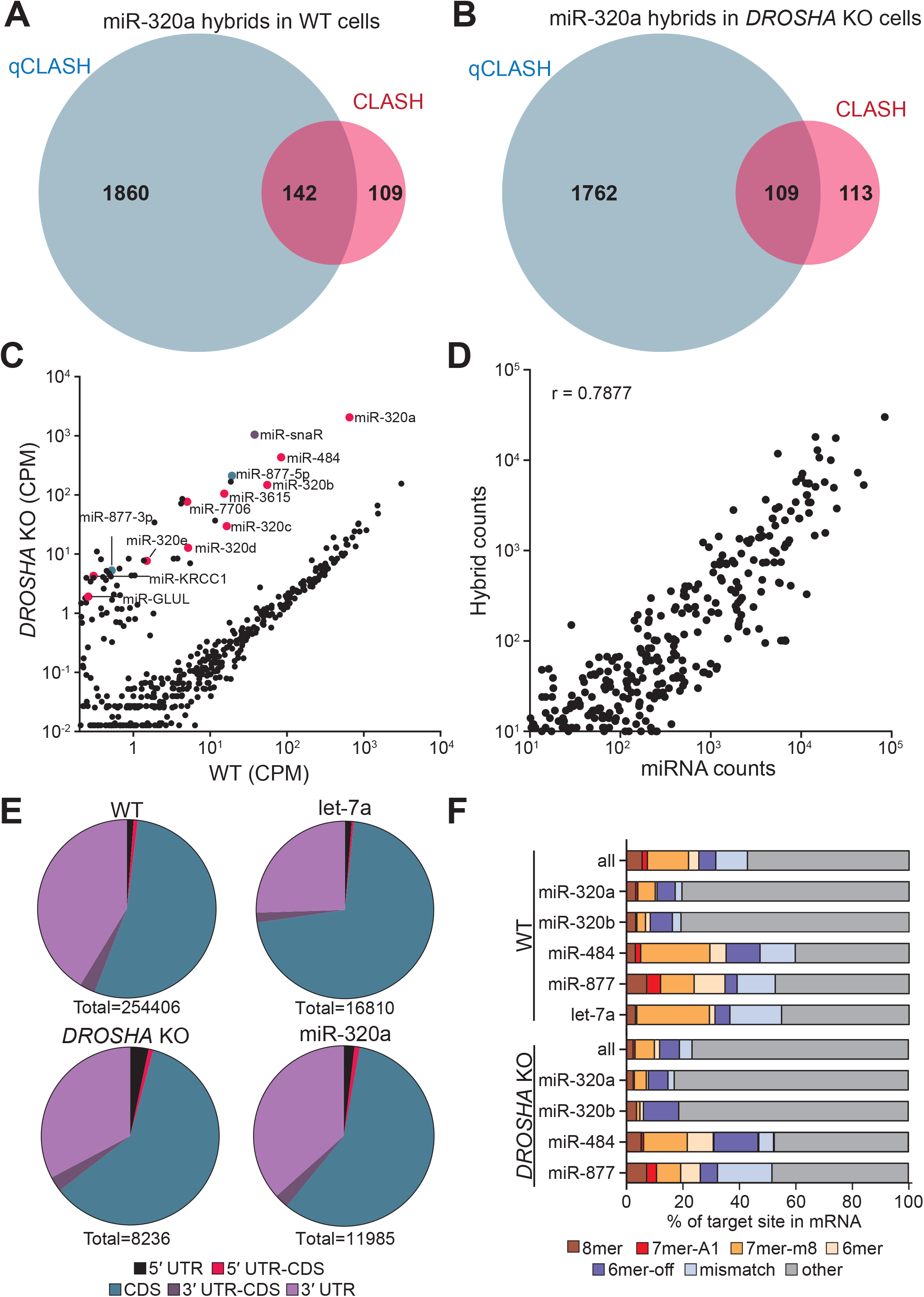
Analysis of AGO-qCLASH in HCT116 colorectal cancer cells. Venn diagram depicting the overlap of miR-320a target genes in qCLASH and CLASH for (A) WT cells and (B) *DROSHA* KO cells. (C) Normalized read count of miRNAs in HCT116 and HCT116 *DROSHA* KO cells. miRNA counts were determined with miR-Deep2 and normalized to total reads. TSS-miRNAs are labeled in magenta, mirtrons are labeled in green, miR-snaR is labeled in purple, and all other miRNAs are labeled in black. (D) Correlation plot of miRNA abundance and hybrid counts for each miRNA. miRNA counts were determined using miR-Deep2 and normalized to total number of reads. The number of hybrids for each miRNA was determined and plotted against miRNA counts. A correlation coefficient (r) is provided. (E) miRNA target sites are found predominately in the CDS and 3′UTR. The location of each transcript from the mRNA portion of the hybrid was determined using data from Ensemble Biomart. Each hybrid was assigned a location based on where it was located in relation to the beginning and end of the reference CDS. Transcripts with IDs not found in the database were excluded from the analysis. (F) The proportion of hybrids with seed binding was less than non-canonical pairings. Seed binding was determined using viennad, which predicts folding between the miRNA and its target. Individual miRNAs display distinct distributions of seed bindings.

Aside from dominant miR-320a hybrids, we found limited numbers of hybrids for other Drosha-independent miRNAs. A possible remedy to identify more targets for less abundant miRNAs is using a miRNA specific primer to generate a qCLASH library (Fig. S2A) (see Materials and Methods for details) (49). As a proof-of-concept experiment, we performed miR-320a-qCLASH using the same cDNA generated from qCLASH. From approximately 1/10 of the raw reads compared with qCLASH, we identified 1,271 miR-320a/mRNA hybrids, a third of what was identified by qCLASH (Table S2). Over half of the miR-320a-qCLASH targets overlapped with the qCLASH targets (Fig. S2B), suggesting that miR-qCLASH is a possible alternative to detect miRNA-specific hybrids. However, given the abundance of miR-320a in HCT116 cells, it is likely to be the only TSS-miRNA having an impactful regulation on gene expression (7). Because of the far greater number of hybrids identified by qCLASH and the high degree of overlap with CLASH and miR-qCLASH hybrids, we opted to perform additional qCLASH replicates moving forward.

### Qualitative analyses of qCLASH data sets

We generated four additional libraries for WT cells and three for *DROSHA* KO and *DICER* KO cells. Consistent with the first qCLASH biological replicate, TSS-miRNA-containing hybrids are enriched in *DROSHA* KO cells but depleted in *DICER* KO cells (Table 1 and Table S2). Comparing the miRNA counts in WT and *DROSHA* KO cells, we found that Drosha-independent miRNAs were elevated in the *DROSHA* KO cells while canonical miRNAs were repressed (Figs. 2C, S3A, and S3B). We were able to identify 23,723 miR-320a/mRNA hybrids between the WT and *DROSHA* KO cells, making up 5.23% and 65.05% of the WT and *DROSHA* KO miRNA/mRNA hybrid reads, respectively. Somewhat surprisingly, even though the miR-320a counts are elevated in *DROSHA* KO compared with WT qCLASH, numbers of miR-320a/mRNA hybrids in both qCLASH data sets are comparable. This is most likely due to consistently lower levels of AGO immunoprecipitated from UV-crosslinked *DROSHA* KO cells for unknown reasons (Fig. S1B, western blot). Amongst all the Drosha-independent miRNAs, miR-320a hybrids were the most abundant, reflecting miR-320a’s dominance in HCT116 cells. Few hybrids could be identified for other TSS-miRNAs (Table S3). In conclusion, we used *DROSHA* KO cells to enrich Drosha-independent miRNAs and identified prevailing miR-320a hybrids in qCLASH.

**Table 1:** Summary of AGO-qCLASH in HCT116 cells. The table depicts the total number of miRNA/mRNA reads in each qCLASH sample. miR-320a/mRNA reads are broken down by replicate and summed for each sample group. Percentage depicts the fraction of miR-320a hybrids in total miRNA/mRNA hybrids. BR = Biological replicate.

|  | HCT116<br>Cell Line | AGO<br>Antibody | miRNA/mRNA<br>Hybrids | miR-320a/mRNA<br>Hybrids |  |
| --- | --- | --- | --- | --- | --- |
| qCLASH | **Wild Type** |  | **303,980** | **15,902** | **5.23%** |
|  | BR1 | 2A8 | 70,124 | 4,511 | 6.43% |
|  | BR2 | 4F9 | 64,561 | 3,695 | 5.72% |
|  | BR3 | 2A8 | 32,940 | 1,942 | 5.90% |
|  | BR4 | 2A8 | 59,250 | 2,399 | 4.05% |
|  | BR5 | 4F9 | 77,105 | 3,355 | 4.35% |
|  | **DROSHA KO** |  | **12,023** | **7,821** | **65.05%** |
|  | BR1 | 2A8 | 4,752 | 3,544 | 74.58% |
|  | BR2 | 2A8 | 1,739 | 1,171 | 67.34% |
|  | BR3 | 4F9 | 2,562 | 1,481 | 57.81% |
|  | BR4 | 4F9 | 2,970 | 1,625 | 54.71% |
|  | **DICER KO** |  | **3,044** | **108** | **3.55%** |
|  | BR1 | 2A8 | 662 | 22 | 3.32% |
|  | BR2 | 2A8 | 1,005 | 42 | 4.18% |
|  | BR3 | 4F9 | 886 | 31 | 3.50% |
|  | BR4 | 4F9 | 491 | 13 | 2.65% |

Previous CLASH studies reported that miRNA abundance correlates with the number of hybrids of each miRNA (34). Averaging five WT qCLASH data sets, we found a strong correlation between miRNA abundance and the number of hybrids identified for each miRNA (r = 0.7877, Fig. 2D). In contrast, we found no positive correlation between transcript abundance and hybrid count (r = −0.04855, Fig. S3C). Therefore, targets enriched in miR-320a hybrids are not due to their transcript abundance, but because of their specific interaction with miR-320a.

We next analyzed the binding location of miRNAs using the qCLASH data and found that miRNAs primarily target the CDS and 3′ UTR as previously reported (Fig. 2E) (30, 33–36). Canonical miRNA interactions involve the miRNA seed region, usually nucleotides 2-7 and sometimes nucleotide 8 on the miRNA (7). However, previous CLASH studies have demonstrated substantial non-seed targeting (33–35). Similarly, we found that non-seed binding, including mismatches, makes up most of the miRNA-target interactions in WT cells (Fig. 2F). Interestingly, we found that miR-320a has more non-seed matches than other miRNAs examined. Given miR-320a’s abundance in *DROSHA* KO cells, it is not surprising that the total miRNA-interactions in *DROSHA* KO closely match miR-320a’s interactions (Figs. 2E and 2F). Taken together, qualitative analyses of the qCLASH data are consistent with previous reports, suggesting that we successfully captured endogenous miRNA/mRNA interactions (33).

### Identification of miR-320a targets with qCLASH

While *DROSH*A KO cells were enriched in Drosha-independent miRNAs including TSS-miRNAs, only miR-320a hybrids were abundant enough to facilitate robust analysis of its functions. To understand the function of miR-320a, we interrogated qCLASH identified targets in at least two replicates in WT or *DROSHA* KO with Ingenuity Pathway Analysis (IPA) to identify cellular pathways regulated by miR-320a (Tables S4 and S5). eIF2 signaling is identified as the highest enriched pathway in WT qCLASH (Figs. 3A & S4A). The top ten most significantly enriched pathways for miR-320a targets in WT qCLASH are also significantly enriched in *DROSHA* KO qCLASH (Fig. 3A).

**Figure 3:**
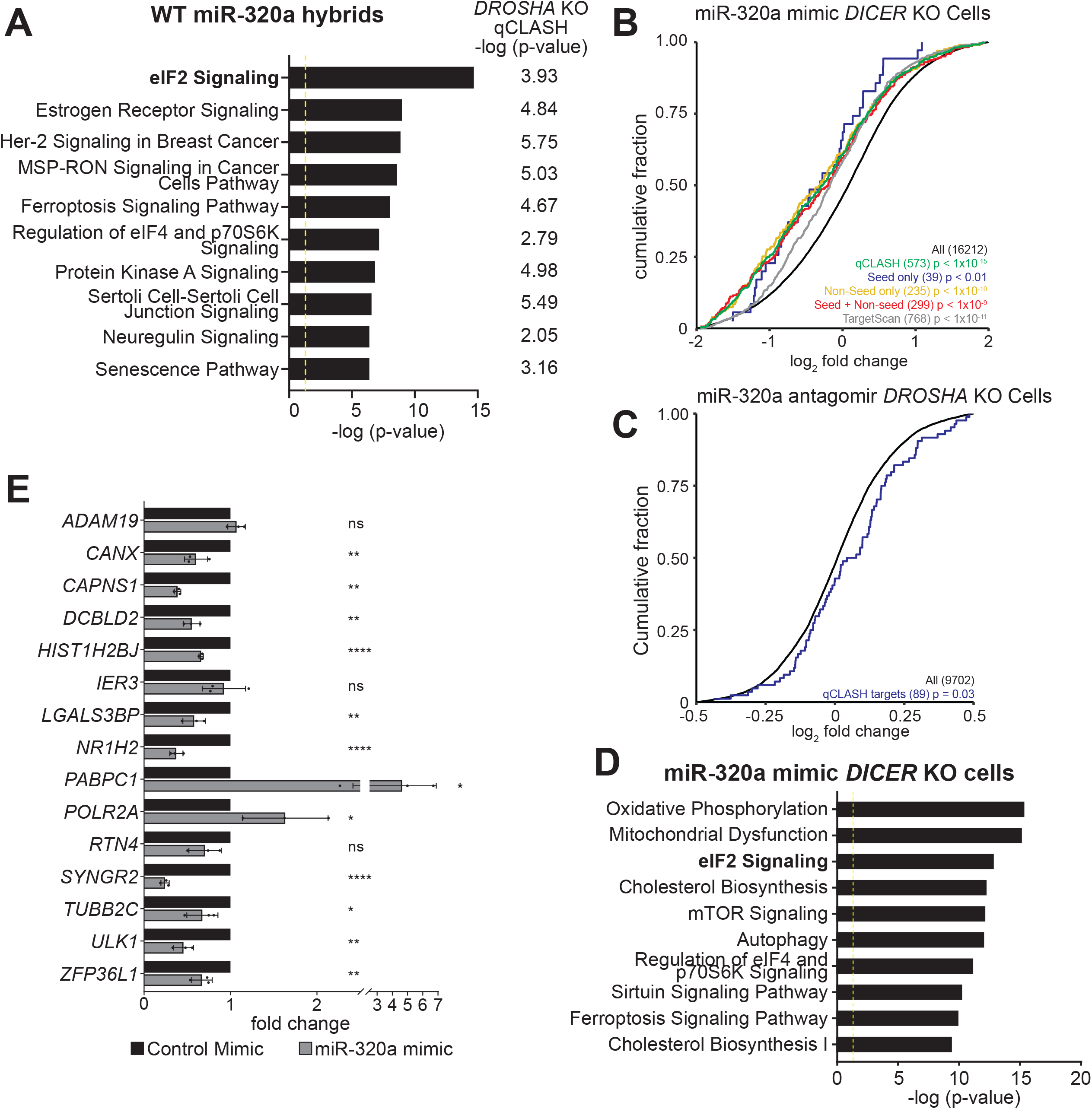
Identification of pathways targeted by miR-320a in HCT116 cells. (A) Top ten pathways targeted by miR-320a in WT qCLASH. The corresponding -log (p-value) for each pathway in *DROSHA* KO qCLASH is depicted right of each bar. The yellow line defines significance threshold for IPA. (B) The cumulative distribution function of fold change in gene expression for qCLASH-identified miR-320a targets (in ≥ 3 WT replicates) following miR-320a mimic transfection in HCT116 *DICER* KO cells. Green line: all qCLASH identified targets; Blue line: qCLASH-identified targets containing only seed matches; Yellow line: qCLASH-identified targets containing only non-seed matches; Red line: miR-qCLASH-identified targets that contain both seed and non-seed matches; Gray line: conserved miR-320a targets predicted by TargetScan; Black line: all genes. P-values were determined using Kolmogorov Smirnov tests between colored subsets and all genes (black). (C) The cumulative distribution function of fold change in gene expression for qCLASH-identified miR-320a targets (in ≥ 4 WT replicates, blue line) following miR-320a antagomir transfection in HCT116 *DROSHA* KO cells. (D) Top 10 pathways enriched in differentially expressed genes (p ≤ 0.05) from (B) were determined by IPA. (E) High-confidence targets from qCLASH were validated using RT-qPCR. HCT116 *DICER* KO cells were transfected with miR-320a mimic or non-specific control. Total RNA was prepared and gene expression was measured with RT-qPCR using the comparative Ct method and depicted as fold changes. Data are shown as the average of three biological replicates with a standard deviation. ns = P > 0.05, * = P ≤ 0.05, ** = P ≤ 0.01, *** = P ≤ 0.001.

Previously, others have demonstrated that predicted miRNA targetomes can be evaluated by measuring the transcriptome-wide gene expression change following exogenous expression of miRNAs (50, 51). Therefore, we used miRNA mimic or antagomir transfection coupled with RNA-sequencing to validate the large number of miR-320a targets identified in qCLASH. It has been noted that transfection of excess miRNA mimics could perturb normal miRNA function via competition with the endogenous miRNAs (22, 52). Since gene expression in *DICER* KO cells is under minimal regulation by endogenous miRNAs, we transfected miR-320a mimic in both WT and *DICER* KO cells and measured global gene expression change using RNA-sequencing. Conversely, antagomir was transfected into *DROSHA* KO cells, which exhibit high endogenous miR-320a expression, to inhibit miR-320a (20). As a validation of the functionality of miR-320a mimic and antagomir, we also transfected WT, *DROSHA* KO and *DICER* KO cells stably expressing a GFP reporter with two complementary miR-320a binding sites (Fig. S5A – upper left panel) (20). In mimic-transfected WT and *DICER* KO cells, a reduction of GFP was observed, signifying that the mimic was loaded onto the RISC complex and is functional (Fig. S5A). In contrast, an increase in GFP was observed in miR-320a antagomir-transfected *DROSHA* KO cells. On northern blots, the mimic drastically increased miR-320a levels in WT and *DICER* KO cells, while an almost complete inhibition of miR-320a was detected in *DROSHA* KO cells transfected with the antagomir (Fig. S5B) (53, 54). These results confirm the efficacy of the miR-320a mimic and antagomir in HCT116 cells.

Consistent with miRNA’s repressive function in mRNA abundance, in RNA-seq of miR-320a-transfected *DICER* KO cells, we found significant repression of qCLASH-identified targets that appear in at least three WT qCLASH replicates (p < 1.0×10^-5^, Fig. 3B, green curve). The degree of repression for qCLASH-identified targets is greater when compared with TargetScan-predicted targets (p < 10^-11^, Fig.3B compare green curve with the gray curve). Given that qCLASH detected a large portion of non-seed targets for miR-320a (Fig. 2F), we examine whether these non-seed targets react differently to miR-320a mimic transfection compared to the canonical seed-match targets. To this end, qCLASH-identified miR-320a targets were classified into three different groups for analysis: targets with only seed match sites (blue curve); targets with only non-seed binding sites (yellow curve), and targets with both seed and non-seed binding sites (red curve). Targets in different groups were repressed by miR-320a mimic to similar extents, suggesting that both seed and non-seed sites are functional (Fig. 3B).

To confirm that miR-320a represses its targets at the physiological level, we analyzed fold changes of miR-320a targets in *DROSHA* KO cells transfected with miR-320a antagomir. miR-320a targets identified in at least three WT qCLASH replicates appeared to be de-repressed, although the change was not statistically significant (p = 0.09, Fig. S5C, red curve). Similarly, TargetScan-predicted targets did not show de-repression compared with all genes (p = 0.14, Fig. S5C, cyan curve). We reasoned that the lack of significant de-repression is because blocking miRNA induces more subtle changes in target expression compared with mimic transfection. Therefore, we further extracted high-confidence miR-320a targets identified in 4 out of 5 WT qCLASH replicates. These targets were indeed significantly de-repressed when miR-320a was inhibited by antagomir (p = 0.03, Fig. 3C), confirming that AGO-qCLASH identified targets are regulated by miR-320 in physiological conditions.

By IPA, the differentially expressed mRNAs in *DICER* KO cells transfected with miR-320a were further analyzed and eIF2 signaling pathway was once again one of the most significantly enriched pathways (p ≤ 0.05) (Figs. 3D and S4B), consistent with the observation that this pathway was enriched in qCLASH-identified targets (Figs. 3A and S4A). Consistently, transcriptomic analyses of miR-320a mimic-transfected WT cells and antagomir-transfected *DROSHA* KO cells show that the eIF2 signaling pathway ranked near the top of the list of most significantly altered pathways determined by IPA (Fig. S6).

To validate individual qCLASH targets, we performed real-time quantitative PCR following reverse transcription (RT-qPCR) on total RNAs extracted from miR-320a-transfected *DICER* KO cells. We selected individual qCLASH targets that appear in 4 of the 5 WT datasets, as well as in 3 out of 4 *DROSHA* KO datasets, generating a list of 15 high-confidence miR-320a targets (Fig. S7). The mRNA portion of each hybrid contains putative miR-320a binding sites that reside within the 3′ UTR or the CDS. Seven of the genes contain canonical miR-320a binding sites in the 3′ UTR. In RT-qPCR, ten of the targets were repressed, two were up-regulated, and three showed no significant change (Fig. 3E). For the targets containing canonical seed matches, all but one was repressed. Therefore, both global RNA-seq and targeted RT-qPCR experiments confirmed that qCLASH identified genuine miR-320a targets.

### miR-320a represses chaperone protein Calnexin

Given that the eIF2 signaling pathway is enriched in miR-320a qCLASH targets and in differentially expressed genes when miR-320a level change, we reasoned that a miR-320a target affecting this pathway would be present in both WT and *DROSHA* KO qCLASH. Among the top 15 high confidence miR-320a targets, the transmembrane endoplasmic reticulum (ER) chaperone protein Calnexin (CANX) directly connects with eIF2 signaling. CANX works in tandem with calreticulin (CALR) to assist in folding glycoproteins or designating unfolded proteins to the proteasome (55–57). When unfolded proteins accumulate in the ER, the three UPR pathways, one of which is the eIF2-mediated ISR, get activated to increase *CANX*/*CALR* expression to help alleviate the stress (58–60). In both RNA-seq and RT-qPCR data, *CANX* mRNA was repressed in response to miR-320a mimic transfection (Figs. 3E and 4A). miR-320a not only lowered *CANX* mRNA, but also reduced the quantity of CANX protein, as western blot-detected CANX protein levels are significantly repressed in WT cells treated with miR-320a mimic (Fig. 4B). When miR-320a was repressed by antagomir in WT cells, upregulation of CANX was observed (Fig. 4C).

**Figure 4:**
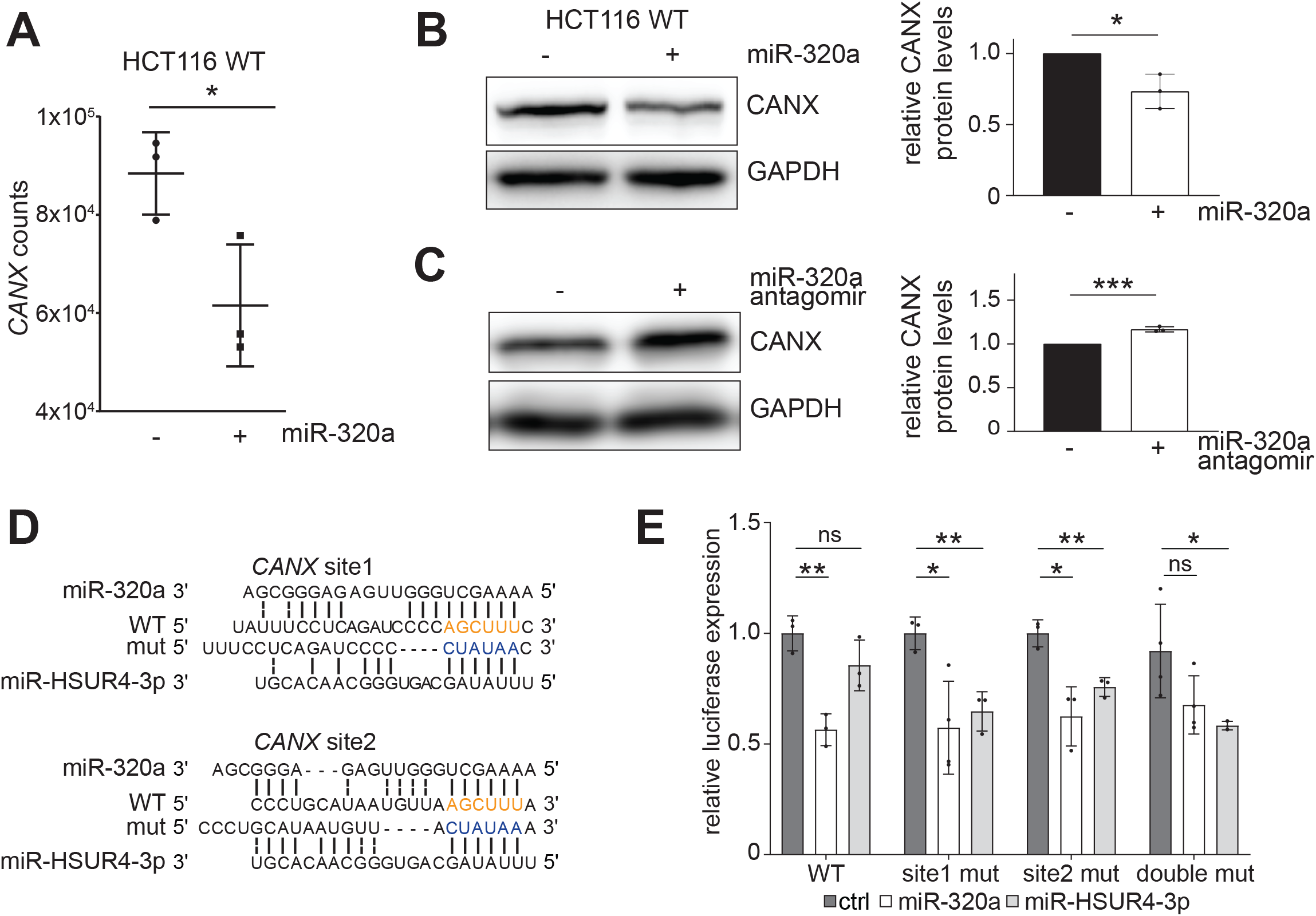
miR-320a targets CANX in colorectal cancer cells to activate UPR. (A) *CANX* expression is reduced in miR-320a mimic transfected *DICER* KO cells, measured by RNA-seq. Gene counts were generated from Stringtie output and normalized using Deseq2. Individual samples were plotted along with mean and standard deviation, for three biological replicates. Western blots of CANX in miR-320a mimic and antagomir (C) transfected HCT116 cells. GAPDH was used as a loading control. Quantitation of CANX protein expression is depicted in a bar graph as individual points with standard deviation for three biological replicates. (D) Predicted base-pairing of miR-320a and miR-HSUR4-3p with *CANX* 3′ UTR; miR-320a seed match sequence on *CANX* is highlighted in orange; miR-HSUR4-3 seed match sequence is highlighted in blue. (E) Dual-luciferase reporter containing WT *CANX* 3′ UTR miR-320a binding sites, a mutation in site 1 (site 1 mut), a mutation in site 2 (site 2 mut), or double mutant was co-transfected with a vector expressing miR-320a, miR-HSUR4-3p, or an empty vector in 293T cells. All luciferase assays were performed in technical and biological triplicates.

In our list of high confidence targets, *CANX* has one miR-320a target sequence in the 3′ UTR (Fig. S7). Additionally, when we examined targets that appeared in fewer qCLASH replicates, we identified an additional binding site within 100 nts of the high confidence *CANX* site. Both sites contain strong canonical seed binding (Fig. 4D). We cloned the *CANX* 3′ UTR containing both qCLASH-identified binding sites downstream of the firefly luciferase within a dual-luciferase reporter. When we co-transfected the WT reporter with a plasmid that encodes miR-320a in HEK 293T cells, reduced expression of the firefly luciferase was observed (Fig. 4E). Using site-directed mutagenesis, we changed both miR-320a binding sites to miR-HSUR4 sites. miR-HSUR4 is a viral miRNA that is not endogenously expressed in 293T cells. In both cases, neither site mutant alone was able to derepress the firefly reporter completely. This finding suggests that a single site is sufficient for miR-320a mediated repression of *CANX*. When both sites are mutated, the firefly luciferase signal increased, demonstrating that both sites are bona fide miR-320a binding sites (Fig. 4E).

We next sought support for miR-320a targeting of *CANX* from patient data available in The Cancer Genome Atlas (TCGA) (61, 62). We found that miR-320a was most down-regulated in colorectal cancer compared to other cancer types (Fig. S8A). Previous reports of miR-320a expression in colorectal cancer demonstrated lower expression when compared to non-tumor tissue (63, 64). Decreasing expression of miR-320a is associated with poor prognosis in colorectal cancer patients (65–67). Conversely, increased *CANX* expression was associated with poor clinical outcomes in colorectal cancer patients (68). We analyzed patient samples where both mRNA and miRNA sequencing was performed using the same tissue (69, 70). miR-320a was down-regulated in six out of the seven tumors, matching previous reports and TCGA data (Fig. S8B) (65–67, 71–73). Conversely, *CANX* was up-regulated in six out of the seven of the tumors analyzed (Fig. S8C). Together with our biochemical characterization of the miR-320a target sites in *CANX* mRNA, these results suggest an inverse relationship between the expression of miR-320a and *CANX*.

### miR-320a activates the unfolded protein response

The accumulation of unfolded proteins in the ER results in the activation of a trio of ER-resident proteins (IRE1, PERK, and ATF6) that triggers three separate signaling pathways, collectively known as the UPR. These three proteins create a torrent of signals directed at either reducing the stress or leading towards programmed cell death (apoptosis) (59). We hypothesized that down-regulation of *CANX* by miR-320a would lead to the accumulation of unfolded proteins. In response to unfolded protein accumulation, IRE1 dimerizes and autophosphorylates, activating its RNase domain to cleave a 26-base unconventional intron out of *XBP1* mRNA (60). This unique splicing allows the translation of the active form of XBP1 which in turn activates the transcription of UPR responsive genes. To determine if miR-320a affects the UPR, we measured the active (i.e., spliced) form of *XBP1* mRNA using RT-qPCR in miR-320a mimic-transfected WT cells. Primers spanning the 26 bp region spliced out by IRE1 were used to measure the amount of spliced *XBP1* (74). We found that miR-320a increases the splicing of *XBP1* (Fig. 5A), in agreement with a previous report that knockout of *CANX* induced IRE1-mediated splicing of *XBP1* and the constitutive activation of the UPR (57). Furthermore, we measured the levels of *BiP* and *CHOP*, downstream genes activated by the UPR. *BiP* and *CHOP* mRNAs in WT cells are upregulated by miR-320a mimic transfection, further supporting the role of miR-320a in activating UPR (Fig. 5A).

**Figure 5:**
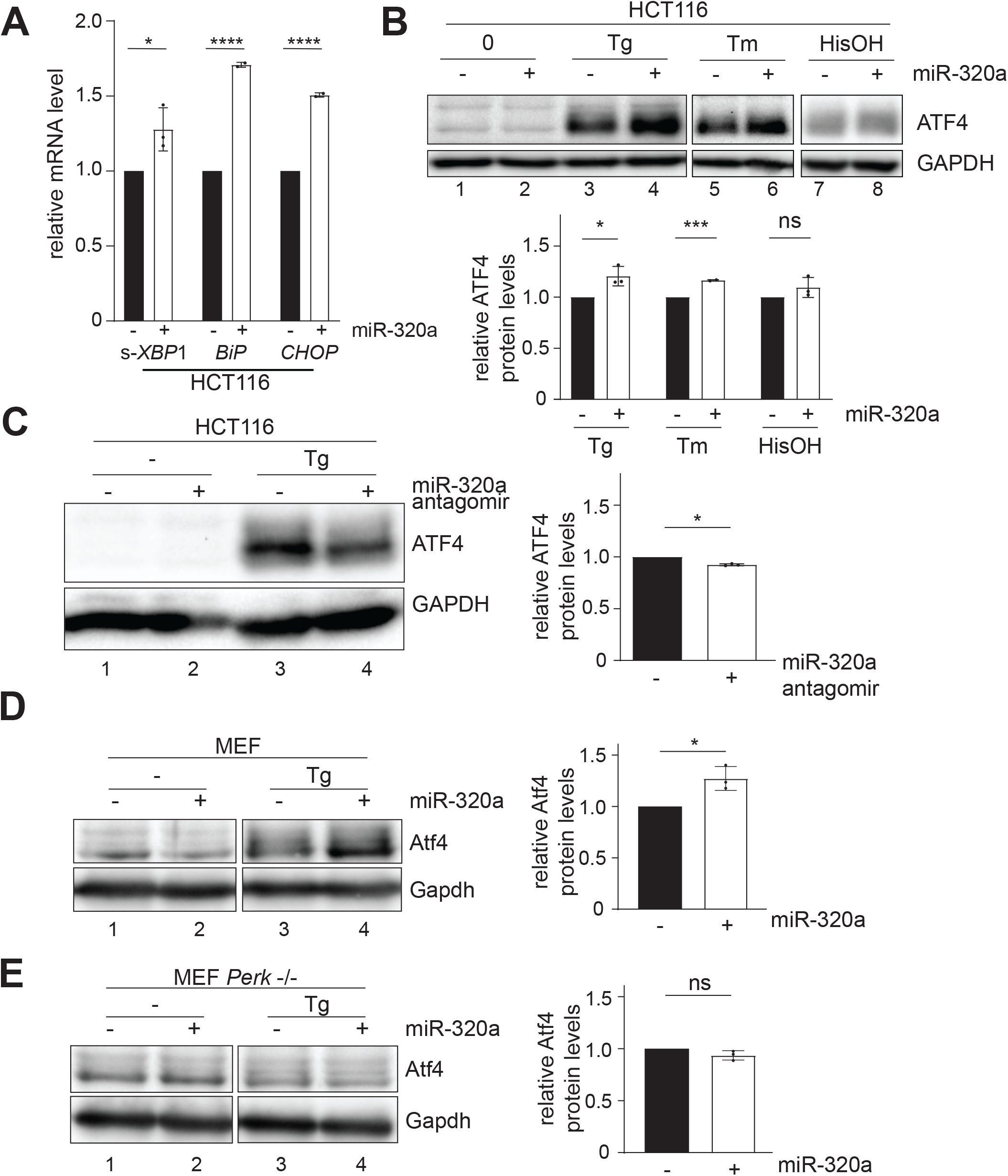
miR-320a activates ATF4 during ER stress. (A) RT-qPCR quantification of spliced(s)-*XBP1*, *BiP*, and *CHOP* mRNA in HCT116 cells transfected with miR-320a mimic. Data are shown as the average of two - three biological replicates with a standard deviation and individual data points. s-*XBP1* was normalized to *HPRT*. Experiments were performed in technical triplicate. (Student’s t-test): ns = P > 0.05, * = P ≤ 0.05, **** = P ≤ 0.0001. (B) Western blots of ATF4 in miR-320a mimic-transfected WT cells with thapsigargin (Tg), tunicamycin (Tm) or histidinol (HisOH) treatment. (C) Western blots of ATF4 in miR-320a antagomir-transfected HCT116 with thapsigargin (Tg) treatment. Western blots of Atf4 in miR-320a mimic-transfected MEF WT (D) and Perk −/− cells (E) with Tg treatment. In all western blots, GAPDH was used as a loading control. Quantitation of three biological replicates in with mean and standard deviation is shown with individual data points. (Student’s t-test): ns = P > 0.05, * = P ≤ 0.05.

Activation of the *PERK* arm of the UPR leads to a suppression of global protein synthesis, but at the same time translational induction of a well characterized effector, Activating Transcription Factor 4 (*ATF4*) (58). This translational control mechanism, termed the ISR, is also observed for other cellular stress responses, including amino acid deprivation leading to the presence of uncharged tRNAs (59, 75, 76). Given miR-320a’s involvement in UPR via targeting *CANX*, we reasoned that miR-320a might affect *ATF4* expression under ER stress induced by unfolded proteins. To this end, we transfected miR-320a in WT HCT116 cells and treated them with either thapsigargin (Tg) or tunicamycin (Tm). Tg is an endoplasmic reticulum Ca^2+^ ATPase inhibitor that prevents the import of Ca^2+^ into the lumen of the ER (60, 77). Tm blocks protein N-glycosylation (78). Both treatments ultimately result in the accumulation of unfolded proteins. As expected, ATF4 protein levels increased upon Tg- and Tm- treatment (Fig. 5B), and cells transfected with miR-320a showed exacerbated up-regulation of *ATF4* (Fig. 5B, compare lane 3 to lane 4, lane 5 to lane 6), while cells transfected with miR-320a antagomir showed lower *ATF4* up-regulation (Fig. 5C, compare lane 3 to lane 4). To test whether the miR-320a up-regulated *ATF4* expression is specific to the UPR, we repeated miR-320a transfection with the addition of histidinol, an amino acid alcohol that mimics stress induced by amino acid deprivation (79). No significant change in *ATF4* expression with or without miR-320a transfection was detected, which suggests the miR-320a-mediated change in *ATF4* is specific to the UPR (Fig 5B, compare lane 7 to lane 8).

Given that upregulation of *ATF4* in response to stress is transient, we conducted a time-course experiment to observe *ATF4* expression in the presence of Tg-treatment and miR-320a transfection over time. As before, we found that miR-320a increases *ATF4* expression in stress-induced HCT116 and HEK 293T cells, which peaked at 4 hours after Tg-treatment (Figs. S9A and S9B). CANX was initially repressed, before being upregulated towards the later time points in miR-320a overexpressed cells, signifying that CANX expression is activated in response to the ATF4 upregulation (Figs. S9A and S9B). During Tg treatment, we observed that miR-320a levels remain constant in HCT116 cells, but miR-320a cannot be detected by northern blot in 293T cells (Figs. S9C and S9D).

We next examined if expressing miR-320a closer to the physiological level in cells would also enhance *ATF4* up-regulation. Human embryonic kidney 293T cells express low levels of miR-320a and transfection of a plasmid containing miR-320a gene under the control of its native promoter enables miR-320a expression (18). Similar to miR-320a mimic transfection in HCT116 cells, we observed up-regulation of *ATF4* in both Tg- and Tm- treated 293T cells when miR-320a was expressed from the plasmid (Fig. S10A, compare lane 3 to lane 4, lane 5 to lane 6).

Finally, we tested whether UPR regulation by miR-320a occurs in other cell lines. Upon transfecting miR-320a mimic in cells with low endogenous miR-320a, such as glioblastoma T98G cells, HeLa cells and mouse embryonic fibroblast (MEF) cells, Tg-induced *ATF4* activation was further elevated (Figs. 5D, S10B and S10C, compare lane 3s to lane 4s). On the other hand, we transfected miR-320a antagomir in medulloblastoma DAOY cells, in which endogenous miR-320a levels are high and treated the cells with Tg. As expected, inhibition of miR-320a dampened the activation of *ATF4* in DAOY cells (Fig. S10D, compare lane 3 to lane 4). With MEF cells, we further utilized a previously established *Perk* −/− line to test whether *Atf4* upregulation is directly connected with UPR (Jixiu Shan and Michael Kilberg, manuscript in preparation). Accumulation of unfolded proteins in the ER causes the dissociation of the protein BiP from PERK, allowing it to dimerize. Auto-phosphorylation of PERK dimer is essential to activate the ISR response. Indeed, in MEF cells without Perk, miR-320a mimic transfection failed to enhance *Atf4* activation (Fig. 5E, compare lane 3 to lane 4). In summary, miR-320a’s regulation of UPR is observed in diverse cell lines.

## DISCUSSION

### qCLASH allows for the identification of endogenous miRNA/target pairing

Bioinformatic predictions based on seed binding of miRNA with mRNA 3′ UTR have expanded our understanding of miRNAs’ functions. However, recent studies using CLASH have identified significant miRNA binding events that do not require perfect seed base-pairing as well as miRNA binding to regions other than the 3′ UTR (33–36, 80). This study identified seven high-confidence non-canonical miR-320a targets including eleven different target sites in both the 3′ UTR and the CDS. Amongst these genes, four were repressed, two were up-regulated, and one was not significantly affected (Figs. 3E and S6B). Likewise, other studies have documented non-canonical pairings. For example, miR-20a exclusively targets *DAPK3* in the CDS region, but not when the site is moved to the 3′ UTR of a luciferase reporter (81). Additionally, miR-20a targeting of *DAPK3* represses gene expression by blocking translation instead of targeting the mRNA for decay. In addition to novel binding sites, several studies have documented the presence and efficacy of non-seed matches, including uridylated miR-124 having a distinct targetome from canonical miR-124 (49, 50, 82). Among the high confidence miR-320a targets we identified here, seed-matched targets in the 3′ UTR were the most robustly suppressed upon miR-320a transfection (Figs. 3E and S6B). The notable exception was *ADAM19*, which was only slightly repressed. Given the abundance of non-canonical sites in our study and many others, they likely represent different paradigms for targeting and function (33–35, 81, 83), while non-repressed targets may represent transient binding events that do not negatively impact target protein expression (32).

Alternative cleavage by Drosha and Dicer and modification by exonucleases and nucleotidyl transferases can produce various templated and non-templated isomiRs (84, 85). IsomiRs can exert different targeting modes, increasing the complexity of miRNA target identification. In particular, 3′ uridylation allows miRNAs to target non-canonical targets without seed matches (86). Since qCLASH ligates miRNAs to their targets, endogenous modifications, like trimming and tailing, could be captured while miRNAs are base-paired with a specific set of targets. The use of qCLASH allows genome-wide identification of such events that would otherwise be passed over with bioinformatic predictions or AGO-CLIP. Additionally, qCLASH can be used to identify novel miRNA with TOMiD (target oriented miRNA discovery), a database-naive bioinformatic method for identifying miRNA in chimeric reads (48). In TOMiD, reads that do not fully align to a single reference sequence are parsed for characteristics of miRNA hybrids. Thus, hybrids for miRNA reads can be identified without reference sequences for miRNAs, allowing a non-biased search for novel miRNAs. Here we presented the traditional utility of qCLASH to identify TSS-miRNA targets.

### miR-320a stimulates the unfolded protein response

TSS-miRNAs are generated in an evolutionarily conserved mechanism that indicates the importance of their expression (19, 20). However, the targets and the roles of TSS-miRNAs remain poorly understood, especially in the context of cancer. Mutations in miRNA processing genes change the miRNA interactome in Wilms tumor, melanoma, and breast cancer (23–25, 27). Furthermore, in breast cancer and Wilms tumor, *DROSHA* down-regulation was coupled with increased expression of Drosha-independent miRNAs (23, 29). Identifying TSS-miRNAs’ targets will be important in understanding the role this class of miRNA have in cancer where *DROSHA* is down-regulated and will facilitate the development of targeted therapies.

Using qCLASH, we were able to identify the targets of miR-320a, the most conserved and abundant TSS-miRNA in HCT116 colon cancer cells (18, 37). qCLASH successfully identified the endogenous HCT116 miRNA targetome, including targets predicted by TargetScan (32). miR-320a is the most abundant TSS-miRNA in WT and *DROSHA* KO cells (Fig. 2C, Table S3). The pathway analysis of qCLASH-identified targets revealed that miR-320a regulates the eIF2 signaling pathway (Fig. 3A). Among the high-confidence targets, the ER chaperone CANX is involved in the unfolded protein response, one arm of which involves increased eIF2 signaling to transiently suppress global protein synthesis. We identified two miR-320a binding sites in the *CANX* 3′ UTR using qCLASH and exogenously expressed miR-320a or inhibited endogenous miR-320a affect *CANX* mRNA and protein levels (Figs. 3E, 4A, 4B, and 4C). Mutating either binding site alone did not abate CANX reporter repression, a decrease was observed when both were simultaneously mutated (Figs. 4D and 4E). We further demonstrated that the expression of miR-320a causes constitutive activation of the UPR, in line with a previous report (Fig. 5A) (57). Finally, we found that miR-320a is down-regulated in colorectal cancers compared to normal adjacent tissue in TCGA data (Fig. S8A). In mRNA and miRNA sequencing data from the same patient-derived tumor, miR-320a was down-regulated (Fig. S8B), while *CANX* expression increased (Fig. S8C).

The expression of miR-320a is frequently down-regulated in colorectal cancer cells. (64, 65, 73), and conversely, overexpression of miR-320a reduced migration, invasion, and proliferation of tumor cells (65, 71, 72). During oncogenesis, the progressive loss of miR-320a expression is associated with tumor progression (66). Low miR-320a expression in colorectal cancer has been linked to increased incidences of metastases, the leading cause of mortality in cancer (66, 87). Loss of miR-320a is predictive of outcomes in colorectal cancer patients, with low expression being correlated with decreased survival (67). Following the excision of tumors, miR-320a concentration in plasma increased in patients who had clinical improvement (73). These results strongly support using miR-320a as a biomarker for disease progression and clinal outcome. It was previously reported that miR-320a targets β-catenin, the transcriptional factor for Wnt signaling (71, 88). Frequently up-regulated, Wnt signaling is regarded as one of the main molecular drivers of oncogenesis in colorectal cancer (89). Understanding how the loss of miR-320a expression in colorectal cancer drives oncogenesis will be important in developing targeted therapies.

Our results demonstrate how qCLASH can identify cellular pathways targeted by miRNAs in cancer cells. Specifically, we identified *CANX* as a miR-320a target, which can trigger the UPR. The accumulation of unfolded proteins results in the activation of *ATF4*, one of the main transcriptional effectors for eIF2 signaling. We demonstrated that miR-320a increases ATF4 abundance during unfolded protein-induced ER stress. Given that miR-320a is repressed during the progression of colorectal cancer, the UPR could be dampened, allowing the tumor cells to escape apoptosis that results from sustained ER stress signaling. In conclusion, we have used AGO-qCLASH to dissect the miR-320a targetome in HCT116 colorectal cancer cells and discovered that TSS-miRNA miR-320a targets *CANX* and stimulates the UPR.

## MATERIALS AND METHODS

### Cell culture

HCT116 cells were maintained in McCoy’s 5A media (Cytiva, SH30200FS), and T98G, Hela, mouse embryonic fibroblasts and 293T cells were maintained in Dulbecco’s Modified Eagle’s medium (Sigma-Aldrich, D5796-500ML) at 37°C with 5% CO2. All media was supplemented with 10% fetal bovine serum and 1X penicillin/streptomycin (Gibco, 15070063). For experiments involving thapsigargin (Sigma, T9033), tunicamycin (Sigma, T7765) and histidinol (Sigma, H6647), media was supplemented with 1X GlutaMAX (Gibco, 35050079) and 1X MEM Non-Essential Amino Acids (Gibco, 11140050).

### AGO-CLASH and qCLASH

AGO-CLASH and qCLASH was conducted on HCT116 WT, *DROSHA* KO, and *DICER* KO cells as previously described with modification from Gay et al (35). Cells were grown on 15 cm plates until 80-90% confluent, washed twice in 1X PBS (137 mM NaCl, 2.7 mM KCl, 10 mM Na_2_HPO_4_, 1.8 mM KH_2_PO_4_), and irradiated with 254 nm UV at 0.6 J/cm^2^. Cell pellets were frozen at −80°C until lysis. Pellets were resuspended in cell lysis buffer (50 mM HEPES-KOH, pH 7.5, 150 mM KCl, 2 mM EDTA, 1 mM NaF, 0.5% NP-40, 0.5 mM DTT), lysed for 15 minutes on ice, treated with 10U for RQ1 DNase (Promega, M610A) for 5 minutes at 37°C with shaking, and centrifuged at 21,000xg for 15 minutes at 4°C.

Six mg of Dynabeads Protein G beads (Invitrogen, 10004D) were washed in PBS-T three times and conjugated with 57.6 μg AffiniPure Rabbit Anti-Mouse IgG (Jackson ImmunoResearch, 315-005-008). After washing three times in PBS-T (1X PBS, 0.02% Tween-20), beads were incubated with 4 μg 2A8 anti-AGO antibody. 156 μg of Protein L Magnetic Beads (Pierce, 88850) were washed three times in PBS-T and incubated with 15.6 μg 4F9 anti-AGO antibody. Beads were washed three times in 1X PXL (1X PBS, 0.1% SDS, 0.5% sodium deoxycholate, 0.5% NP-40) and once with lysis buffer. Approximately 3 mg of lysate were pre-cleared with 3 μg unconjugated Protein G beads or 7.8 μg Protein L beads to remove non-specific binding.

The pre-cleared lysate was added to beads and incubated overnight at 4°C with gentle agitation. Lysates were removed and the beads were washed three times in lysis buffer. Each sample was incubated in 15 ng/μL RNaseA for 12 minutes at 22°C. Intermolecular ligation and libraries were generated as previously described (Supplemental Materials and Methods) (35). Libraries were separated on an 8% native polyacrylamide gel and regions between 147 and 527 bps were excised and eluted overnight in 1:1 RNA elution buffer (300mM sodium acetate, 25 mM Tris-HCl pH 8.0): PCA. The excess adapter was removed using AMPure XP beads (Beckman, A63880) following the manufacturer’s instructions. Libraries were sequenced on the Illumina HiSeq 3000 by the University of Florida Interdisciplinary Center for Biotechnology Research (ICBR) NextGen DNA Sequencing Core.

### Bioinformatic analysis of qCLASH libraries

Sequencing reads from qCLASH libraries were first analyzed with Trimmomatic to remove the adapter and low-quality reads (42). PEAR was used to combine forward and reverse reads into a single forward read (43). PCR duplicates were removed using fastx_toolkit and the unique molecular identifier was removed by trimming the first four and last four nucleotides with Cutadapt (45, 46). Finally, reads were analyzed with Hyb to generate hyb and viennad files. A custom script was used to identify clusters between replicates for each miRNA. Individual miRNA counts from adapter-trimmed sequencing data were determined using miR-Deep2 (90). The genome used for mapping was hg38 and miRNA sequences were obtained from miRbase v.22 (91). Previously identified TSS-miRNAs were manually added to the miRbase file (19, 20, 37). miRNA binding and miRNA seed matches were determined using custom python scripts. All scripts used in this study are available at GitHub (https://github.com/UF-Xie-Lab/qCLASH).

### Quantitative Reverse-Transcription Polymerase Chain Reactions (RT-qPCR) and RNA-seq

For RT-qPCR, 1×10^6^ HCT116 *DICER* KO cells were transfected with 10-20 pmols of either negative control or miR-320a mimic with 7.5 μL of Lipofectamine RNAiMAX (Invitrogen, 13778150) in a 6-well plate. After 48 hours, total RNA from cells were extracted using TRIzol (Invitrogen, 15596018). Total RNA was treated with 1U/μg RQ1 DNase at 37°C for 30 minutes. RNA was purified by PCA and resuspended in nuclease-free water.

For RT-qPCR, DNase-treated RNA was reverse transcribed using iScript Reverse Transcription Supermix (BioRad, 1708841). The resulting cDNA was used for qPCR using the SsoAdvanced Universal SYBR Green Supermix (BioRad, 1725275). Each reaction was performed in technical and biological triplicate using 12.5 ng of cDNA. The cycling conditions used were initial denaturation at 95°C for 30 seconds, and 40 cycles of 95°C for 15 seconds and 60°C for 30 seconds. Fold changes were determined using the comparative Ct method (92). Primers used for RT-qPCR are listed in Table S1.

For RNA-seq of mimic and antagomir transfected cells, polyA-RNAs were enriched from total RNA and sequenced by UF ICBR and Novogene. At UF ICBR, libraries were made using the NEBNext Ultra II Directional RNA Library Prep Kit following the polyA workflow (NEB, E7760). A custom protocol was used at Novogene for libraries preparation. Sequencing reads from poly-A selected RNA-seq were processed with Cutadapt to remove the residual adapter and filter for shorts reads using the parameter “-m 17” (46). Trimmed pair-end reads were aligned to the human genome (Gencode - GRCh38.013) using HISAT2 with the parameters “--dta” (93). Samtools was used to convert the output of HISAT2 into a sorted bam file (94). Reads were assembled using StringTie and a reference transcriptome (Gencode – v37 primary Assembly) with the parameters “-e −B” (95). Differential gene expression for assembled transcripts was determined using Ballgown as previously described (96). Counts for genes were generated from the StringTie output using the supplied script prepDE.py3. DEseq2 was used to generate normalized counts (97).

For gene expression of *DICER* KO cells, total RNA was extracted and polyA selected libraries were prepared by Novogene. Data analysis was conducted by the UF ICBR Bioinformatics Core. Briefly, Trimmomatic was used to remove adapter sequences and poor quality reads (42). Reads were aligned to the genome (GRCh38 - https://useast.ensembl.org/Homo_sapiens/Info/Index/) using STAR (98). Gene expression was determined by extracting the FPKM from aligned reads using RSEM (99).

### Pathway analysis

Ingenuity Pathway Analysis (IPA) (Qiagen) was used to identify pathways targeted by miR-320a. We identified 815 miR-320a targets that appear in two out of five replicates from WT cells and 327 targets that appear in two out of four replicates from *DROSHA* KO cells. Each set was run through IPA’s core analysis to identify significantly targeted pathways. For RNA-seq experiments, differentially expressed genes (p ≤ 0.05) were analyzed using IPA’s core analysis.

### Antibodies and Western blot

HCT116 and 293T cells were sonicated in RIPA lysis buffer (50 nM Tris-HCL pH 8.0, 1% Triton X-100, 0.5% sodium deoxycholate, 0.1% SDS, 150 nM sodium chloride, 2 mM EDTA pH 8.0, 1X cOmplete EDTA-free (Roche, 05056489001)) until cleared. Protein concentration was determined using the DC Protein Assay Kit II (BioRad, 5000112) following the manufacturer’s instructions. Lysates were separated with sodium dodecyl sulfate polyacrylamide gel electrophoresis (SDS-PAGE 8-10%) and transferred to 0.2 μm nitrocellulose for western blot. Primary antibodies were ɑ-AGO2 antibody 11A9, GAPDH (1:5000-1:10000; Cell Signaling, 14C10), ATF4 (1:5000; Cocalico Biotechnology Inc.), and CANX (1:1000; Cusabio, CSB-PA00899A0Rb) and secondary antibodies were goat anti-rat IgG, HRP conjugated (Pierce, 31470) and goat anti-rabbit IgG, HRP conjugated (Pierce, 31460).

### Northern blot

Northern blots were performed as previously described with either near infrared probe or 32P probe (20, 53, 54, 100). Probes are listed in Table S1. Total RNA was separated on 15% SDS-PAGE and transferred to nitrocellulose. UV crosslinking was performed for HCT116 WT and HCT116 *DICER* KO total RNA treated with mimic. EDC crosslinking was performed for HCT116 *DROSHA* KO total RNA.

### Integrated stress response induction

Cells were seeded in 6-well and 12-well plates at a density of 1.25×10^5^-5×10^5^ cells per well and transfected with miR-320a mimic (Qiagen, 339173 YM00471432) or antagomir (GenePharma) 18-24 hours after seeding with Lipofectamine RNAiMax. A plasmid containing endogenous miR-320a gene was transfected with PEI (18). The media was replenished 12 hours after transfection. 50 nM thapsigargin, 2 μg/mL tunicamycin, or 2 mM histidinol was added 24 hours after transfection and incubated for the designated time before being collected for western blot.

### Luciferase assays

Regions corresponding to the qCLASH-identified miRNA binding site along with flanking regions were PCR amplified using Phusion High-Fidelity DNA Polymerase. CANX fragments were cloned into pmirGlo (Promega, E1330) using In-Fusion HD Cloning Kit (Takara, 639649). Mutants for miR-320a sites were generated using site-directed mutagenesis by replacing the miR-320a seed match sequence with the HVS miR-HSUR4 seed match sequence. For luciferase assays, 2.5×10^5^ 293T cells per well were plated in a 12-well plate. After 18-24 hours of incubation, cells were transfected with 500 ng of reporter and miRNA-vector using 4 μg polyethylenimine (PEI) (Polysciences Inc, 24765-1). Dual-luciferase assays were carried out using the Dual-Luciferase Reporter Assay System (Promega, E19080). After 48 hours, cells were lysed with 1X passive lysis buffer for 15 minutes at room temperature with shaking. Firefly luciferase activity was measured by adding 25 μL Luciferase Assay Buffer II (LARII) to 5 μL of lysate and measured immediately with the GloMax 20/20 Luminometer (Promega, E5311). Reactions were stopped and Renilla luciferase activity was measured by adding 25 μL Stop&Go reagent. Samples were measured in technical and biological triplicate.

### miR-320a expression in The Cancer Genome Atlas

Normalized expression of miR-320a in cancerous tissues and adjacent normal tissues from the TCGA Research Network (https://www.cancer.gov/tcga) was acquired from UCSC Xena (https://xenabrowser.net/) (61, 62).

## DATA AVAILABILITY

Data generated in this publication have been deposited to the NCBI Gene Expression Omnibus (GEO) and can be accessed at GSE164634.

## ACKNOWLEDGEMENTS

We thank Drs. Edward Chan, William Dunn, Narry Kim, Michael Kladde, Grzegorz Kudla, and Zissimos Mourelatos for sharing reagents and protocols; UF Interdisciplinary Center for Biotechnology Research for services in high-throughput DNA Sequencing, and AGO 4F9 antibody production; UF Health Cancer Center Biostatistics & Quantitative Sciences Shared Resources for access to Ingenuity Pathway Analysis and the UF Health Cancer Center Cancer Informatics Shared Resource.

## AUTHOR CONTRIBUTIONS

C.F. and M.X. conceived the project. C.F performed AGO-qCLASH and analyzed data with assistance from L.G. and R.R.. L.L. developed scripts for data analyses and identified TCGA data for analysis. T.H. performed miR-320a qCLASH. T.L. performed western blot for UPR genes. N.H. and P.S. performed RT-qPCR for UPR genes. T.G. helped analyze IPA and RNA-seq data. UPR experiments were conducted under the direction of J.S and M.K.. J.B. identified colorectal cancer data sets for analysis. C.F., M.S.K., and M.X. wrote the manuscript.

