## Supplemental Methods and Figures for "Sequencing of Argonaute-bound miRNA/mRNA hybrids reveals regulation of the unfolded protein response by microRNA-320a"

**SUPPLEMENTAL MATRIALS AND METHODS**

**Intermolecular ligation and library preparation for AGO-CLASH and qCLASH**

Following RNase digestion, the beads were washed three times each with 1X PXL, 5X PXL (5X PBS, 0.1% SDS, 0.5% sodium deoxycholate, 0.5% NP-40), High-Stringency Buffer (15 mM Tris-HCL pH 7.5, 5 mM EDTA, 2.5 mM EGTA, 1% Triton X-100, 1% sodium deoxycholate, 0.1% SDS, 120 mM sodium chloride, 25 mM potassium chloride), High Salt Buffer (15 mM Tris-HCl pH 7.5, 5 mM EDTA, 2.5 mM EGTA, 1% Triton X-100, 1% sodium deoxycholate, 0.1% SDS, 1 M sodium chloride), and 1X PNK buffer (50 mM Tris-HCl pH 7.5, 10 mM magnesium chloride, 0.5% NP-40). Following washes, the samples were phosphorylated with 40U T4 PNK (NEB, M0201L) at 16°C for 40 minutes, washed three times with 1X PNK buffer, and intermolecular ligation performed with 500U T4 RNA Ligase I (NEB, M0437M) at 4°C overnight. Beads were washed three times with 1X PNK buffer and dephosphorylated with 3U Calf Intestinal Alkaline Phosphatase (NEB, M0290) at 16°C for 40 minutes. Beads were washed twice with 1X PNK buffer-EGTA (50 mM Tris-HCl pH 7.5, 20 mM EGTA, 0.5% NP-40) and three times with 1X PNK buffer. For qCLASH, eighty pmols of RNA 3′ Adapter (RA3R) was ligated to the RNA with 80U of T4 RNA Ligase 2, truncated K227Q (NEB M0351L) in 1X T4 RNA Ligase Buffer, 10% PEG8000, and 80U of murine RNase Inhibitor (NEB, M0314L) in a total volume of 80 µL at 16°C overnight. CLASH samples were ligated with $^{32}P$ labeled RA3.

After 3′ adapter ligation, CLASH samples were washed three times with 1X PNK buffer and phosphorylated with 40U T4 PNK (NEB, M0201L) in 1X PNK Buffer, 80U of murine RNase Inhibitor, and 37.5 µCl $^{32}P$-ATP at 37°C for 30 minutes with mixing. 80 nmoles of ATP was added and incubate for 15 minutes. The reaction was removed and washed with Hi-Stringency Buffer, Hi-Salt Buffer, 5X PXL Buffer, 1X PXL Buffer, and 1X PNK Buffer. Samples were resuspended in 1X NuPage Buffer, 100mM DTT diluted in PNK buffer and boiled at 75°C for 15 minutes, separated on a 4-12% NuPAGE gel, transferred to a nitrocellulose membrane, and then imaged on a Li-Cor Odyssey. Two regions corresponding to 100-125 and 125-150 kD were excised. qCLASH samples (to be processed without PAGE) were washed three times with 1X PNK buffer and protein/RNA complexes were eluted twice with 100mM sodium bicarbonate and 1% sodium dodecyl sulfate at room temperature for 15 minutes. The two eluates are combined for RNA extraction.

Proteinase K was added to PK buffer (500 mM Tris-HCl pH 7.5, 250 mM sodium chloride, 50 mM EDTA) for a final concentration of 4 mg/mL. RNA was isolated from protein by adding 50 µL (qCLASH) or 160 µL (CLASH) of the diluted proteinase K mix to each sample at 37°C for 20 minutes. Phenol was added to each reaction and incubated at room temperature for 8 minutes with 1400rpm of shaking. The samples were centrifuged at 18,000xg at 4°C for 10 minutes. The aqueous layer (~200 µL) was transferred to a new tube and 20 µL 3M NaOAc, 2 µL 15 mg/mL GlycoBlue (Invitrogen, AM9516), and 500 µL 1:1

mix of ethanol and isopropanol was added to each reaction and then incubated overnight at -20°C or -80°C for 1 hour. Following incubation, samples were centrifuged at 21,000xg at 4°C for 30 minutes and washed two times with 80% ethanol, centrifuging at 18,000xg at 4°C for 10 minutes. Samples were resuspended in nuclease-free water and phosphorylated with 10U of T4 PNK in reaction buffer (1X T4 PNK Ligase Buffer, 20U murine RNase Inhibitor, 15 nmols of ATP) in a volume of 15 μL at 16°C for 40 minutes. Reactions were purified twice by phenol:chloroform:isoamyl alcohol (Fisher, BP17521-400) (PCA), washed twice with chloroform and resuspended in 10 μL water. 100 pmols of RNA 5′ Adapter (RA5R for qCLASH and RA5 for CLASH) was ligated to the RNA with 10U T4 RNA Ligase in reaction buffer (1X T4 RNA Ligase Buffer, 2 μg of BSA, 20U murine RNase Inhibitor, 20 nmols of ATP) at 16°C overnight. The next day, samples were extracted with equal volume of PCA, vortexed at 1400 rpm for 8 minutes at room temperature, and centrifuged at 18,000xg for 10 minutes at room temperature. The aqueous layer (~200 μL) was transferred to a new tube, and 20 μL 3M NaOAc pH 5.2, 30 mg GlycoBlue (Invitrogen, AM9516), and 500 μL 1:1 mix of ethanol and isopropanol was added, and incubated at -80°C for 1 hour. Reactions were centrifuged at 21,000xg for 30 minutes, and the pellet was washed two times with 80% cold ethanol, and centrifuged at 18,000xg for 10 minutes after each wash. The pellet was allowed to dry and resuspended in RNase-free water.

To prepare cDNA libraries, RNA was incubated with 10 pmols each of RTP primer and dNTPs at 65°C for 5 minutes and reverse transcribed with 200U of SuperScript IV Reverse Transcriptase (Lifetech, 18090010) in 1X SuperScript RT Buffer, 100 nM DTT, and 40U murine RNase Inhibitor at 50°C in a reaction volume of 20 μL for 45 minutes, 55°C for 15 minutes, and 95°C for 5 minutes. To determine ideal PCR amplification for each library, 2% of the cDNA was PCR amplified for 10, 12, and 14 cycles with 1U Phusion High-Fidelity DNA Polymerase (NEB, M0530L), 1X Phusion HF buffer, 10 pmols dNTP mix, 2.5 pmols RP1, and 2.5 pmols RPI1. Half the cDNA was used to generate the library using 3 cycles less than what was pre-determined with 0.08U Phusion High-Fidelity DNA Polymerase (NEB, M0530L), 1X Phusion HF buffer, 0.8 pmols dNTP mix, 2 pmols RP1, and 2 pmols RPI.

#### SUPPLEMENTARY FIGURE LEGENDS

**Figure S1: AGO-CLASH library preparation.** (A) AGO immunoprecipitation with 2A8 and 4F9 AGO antibodies in HCT116 cells. Cells were lysed and incubated with antibody-bound beads. I = input, S = supernatant, and P = pellet. (B) RNA enriched by AGO-IP was labeled with $^{32}P$ using T4 PNK, separated on SDS-PAGE, and transferred to nitrocellulose. The boxes mark where the membrane was excised for CLASH library prep. AGO2 was detected by Western blot with the 11A9 antibody. (C) 2% of each cDNA library was PCR amplified with 10, 12, and 14 cycles to determine the optimal number of cycles (3 less than when the smear begins to appear). The boxes indicate the expected library range.

**Figure S2: miRNA-CLASH methodology isolated similar target genes as AGO-CLASH.** (A) Schematic depicting the principle of a miR-specific primer to generate a single miRNA qCLASH library. The RP1 primer has the miR-320a sequence (minus 2 nucleotides) added to the 3′ end. (B) Venn diagram depicting the overlap of miR-320a target genes in qCLASH and miR-qCLASH.

**Figure S3: Qualitative analysis of qCLASH. (**A) *DROSHA* KO results in a global reduction of canonical miRNAs. The expression of each miRNA was normalized to miR320a in *DROSHA* KO cells compared to WT cells. TSS-miRNAs are labeled in magenta, mirtron miR-877 labeled in green, miR-snaR labeled in purple. (B) Drosha-independent miRNAs are ranked high in *DROSHA* KO cells. The expression of miRNAs was determined using miR-Deep2 and averaged together. The top 200 miRNAs expressed in WT cells were compared to their ranking in *DROSHA* KO cells. (C) No correlation was found between the number of miR-320a hybrids identified for each gene in qCLASH and the expression of the gene (RPKM) in *DICER* KO cells. The correlation coefficient (r) is depicted on the graph.

**Figure S4: miR-320a is involved in the eIF2 signaling pathway.** (A) miR-320a targets from WT qCLASH are enriched in IPA analysis. Purple outlines denote that miR-320a targets a component or process, and grey boxes denotes a direct target of miR-320a as determined by qCLASH. (B) Differentially expressed genes in miR-320a transfected *DICER* KO cells are enriched in the eIF2 pathway. Green denotes down-regulation, red denotes upregulation, grey boxes are not significant ($p > 0.05$).

**Figure S5: miR-320a mimic and inhibitor transfection in HCT116 WT, *DROSHA* KO, and *DICER* KO cells.** (A) GFP reporter containing two complementary miR-320a binding sites in the 3′ UTR was expressed in WT, *DROSHA* KO, and *DICER* KO. *Top right* – WT cells were co-transfected with miR-320a and GFP reporter. *Bottom left – DROSHA* KO and *Bottom right – DICER* KO cells stably expressing GFP reporter were transfected with miR-320a antagomir and mimic, respectively. miR-320a mimic suppressed the GFP reporter in *DICER* KO and WT cells. Antagomir suppresses miR-320a expression signified by increased GFP expression in *DROSHA* KO cells. Images were taken 48 hours after transfection. (B) irNorthern blot of miR-320a and U6 in total RNA extracted from WT, *DROSHA* KO, and *DICER* KO cells treated with miR-320a mimic or antagomir. (C) Cumulative distribution of mRNA fold changes in miR-

320a antagomir transfected HCT116 *DROSHA* KO cells; Blue line: conserved miR-320a targets from TargetScan; Red line: qCLASH-identified miR-320a targets appeared in three replicates in WT cells; Black line: all genes. P-values were determined using Kolmogorov Smirnov tests between colored subsets and the all genes control.

**Figure S6: Ingenuity Pathway Analysis (IPA) reveals that miR-320a targets eIF2 signaling.** (A) WT cells were transfected with miR-320a mimic. Differentially expressed genes determined by polyA RNA-seq ($p \leq 0.05$) were analyzed with IPA, where eIF2 signaling is in the top 10 enriched pathways. (B) *DROSHA* KO cells were transfected with miR-320a antagomir. Differentially expressed genes determined by polyA RNA-seq ($p \leq 0.05$) were analyzed with IPA, revealing eIF2 signaling as significantly enriched. The dotted line represents IPA's cutoff for significantly expressed genes( $-\log_2$ p-value $\geq 1.3$).

**Figure S7: Predicted base-pairing interactions in high confidence miR-320a hybrids identified in qCLASH.** High confidence targets with canonical seed matches (A) and non-canonical seed matches (B) are depicted. The microRNA seed region is marked as magenta. The target site's location in the transcript is denoted.

**Figure S8: miR-320a and *CANX* are negatively correlated in colorectal cancer.** (A) The Cancer Genome Atlas (TCGA) gene expression of miR-320a in cancer cells compared to surrounding healthy tissue. Significant changes ($p \leq 0.05$) between Normal (N) and Tumor (T) are marked in red. Patient-derived miRNA and mRNA sequencing data of (B) miR-320a and (C) *CANX* expression in tumor (CRC) compared to adjacent non-tumor cells (ctrl). Significant differences were measured with paired T-test. ns = $P > 0.05$, * = $P \leq 0.05$.

**Figure S9: miR-320a affects UPR/ISR.** Western blot of ATF4, CANX and GAPDH in miR-320a mimic-transfected (A) HCT116 cells and (B) 293T cells treated with thapsigargin (Tg) for various times. irNorthern for miR-320a in HCT116 and 293t cells treated with (C) thapsigargin (Tg) and (D) tunicamycin (Tm). * marks a non-specific band detected by miR-320a probe in 293T cells.

**Figure S10: miR-320a affects UPR in different cell lines.** (A) Representative western blot of ATF4 in miR-320a expression plasmid-transfected HEK293T cells treated with Tg or Tm. Representative western blot of ATF4 in miR-320a mimic-transfected T98G (B), HeLa (C) cells and antagomir-transfected DAOY cells (D) with Tg treatment. GAPDH serves as a loading control. Relative ATF4 levels after miR-320a mimic or antagomir transfection is indicated below the blot.

**Table S1: List of oligonucleotides used in this study.**

**Table S2: Summary of CLASH, qCLASH, and miR-qCLASH.** Summary of CLASH, qCLASH, and miR-320a-qCLASH data. “AB” depicts which AGO antibody was used for IP. “Region” refers to the area on the membrane that was excised for CLASH. Percentage depicts the fraction of miRNA hybrids of the specific miRNA in total miRNA/mRNA hybrids. BR = Biological replicate.

**Table S3: Summary of qCLASH.** Table summary of biological replicates of qCLASH. “AB” depicts which AGO antibody was used for IP. Percentage depicts the fraction of hybrids of the specific miRNA in total miRNA/mRNA hybrids. BR = Biological replicate.

**Table S4: AGO-qCLASH identified miR-320a targets in WT cells.** Table of miR-320a targets in WT cells identified in at least two biological replicates. Replicate refers to the number of different samples each hybrid is identified in. Peak refers to the total number of reads from all replicates.

**Table S5: AGO-qCLASH identified miR-320a targets in *DROSHA* KO cells.** Table of miR-320a targets in *DROSHA* KO cells identified in at least two biological replicates. Replicate refers to the number of biological samples each hybrid is identified in. Peak refers to the total number of reads from all replicates.

### Figure S1

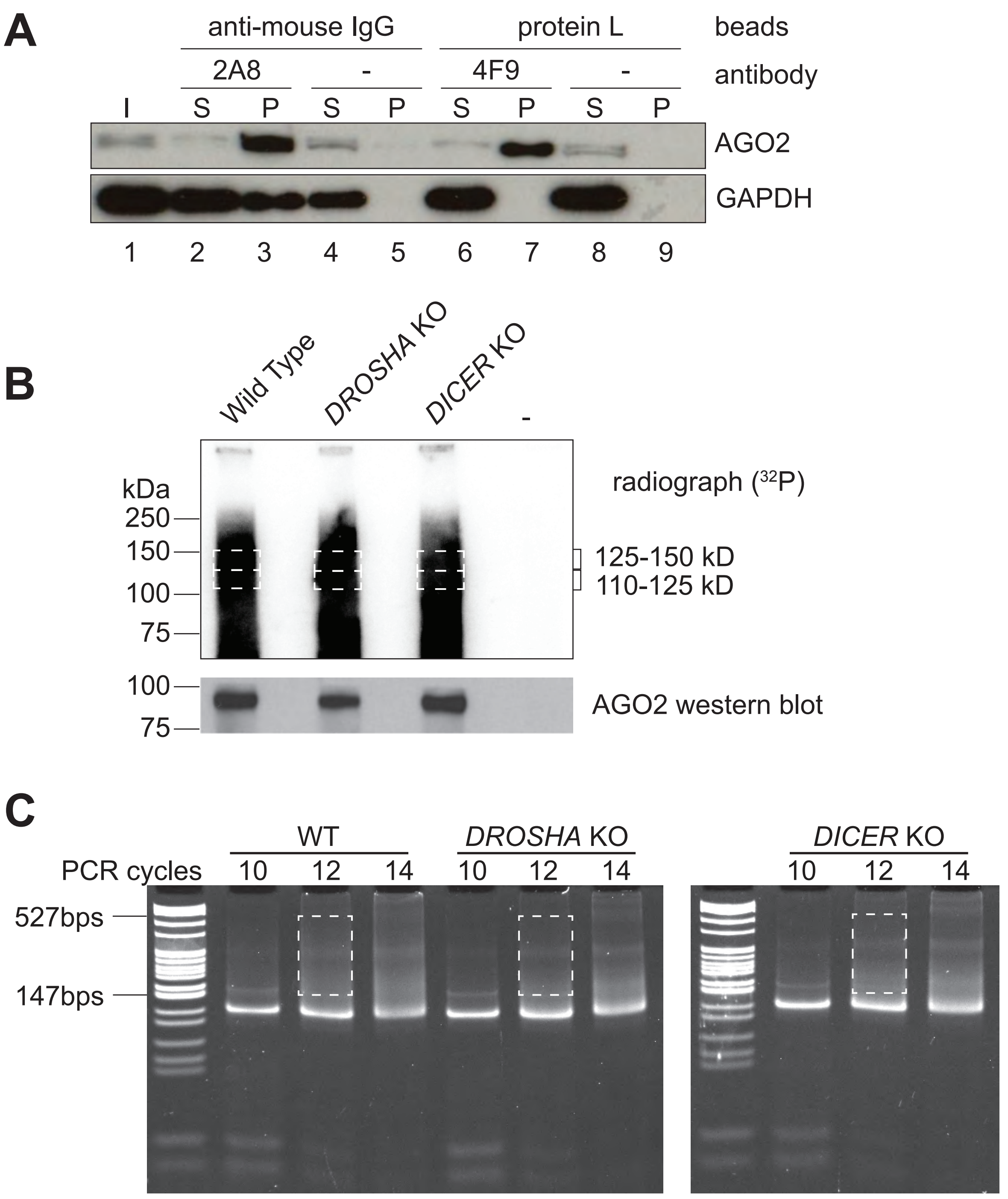

A
anti-mouse IgG
protein L
beads
2A8
-
4F9
-
antibody
I S P S P S P S P
AGO2
GAPDH
1 2 3 4 5 6 7 8 9
B
Wild Type
DROSHA KO
DICER KO
-
kDa
250
150
100
75
radiograph (32P)
125-150 kD
110-125 kD
100
75
AGO2 western blot
C
WT
DROSHA KO
DICER KO
PCR cycles
10 12 14 10 12 14 10 12 14
527bps
147bps

### Figure S2

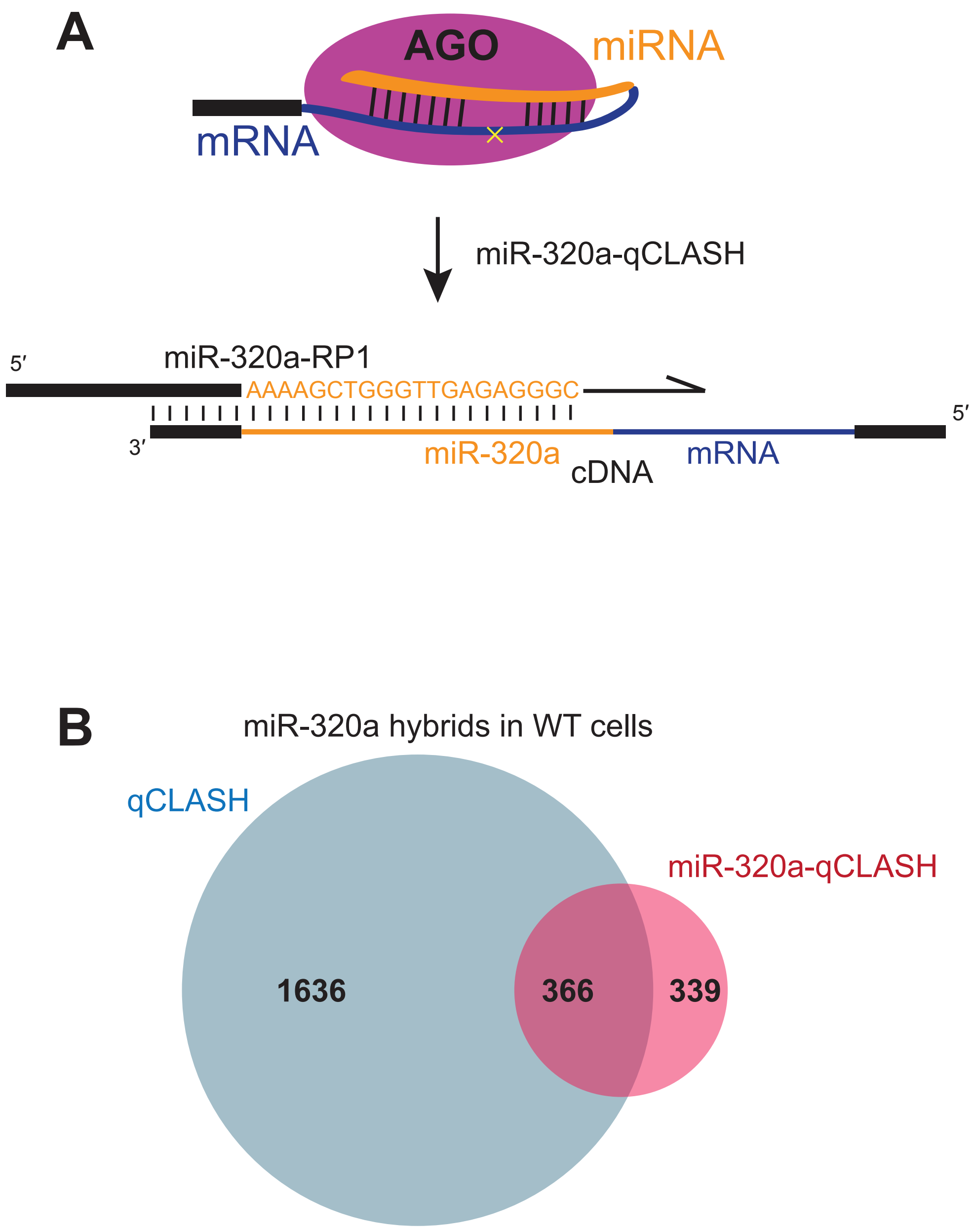

A
AGO
miRNA
mRNA
miR-320a-qCLASH
5′
miR-320a-RP1
AAAAGCTGGGTTGAGAGGGC
3′
5′
miR-320a
cDNA
mRNA
B
miR-320a hybrids in WT cells
qCLASH
miR-320a-qCLASH
1636
366
339

### Figure S3

A

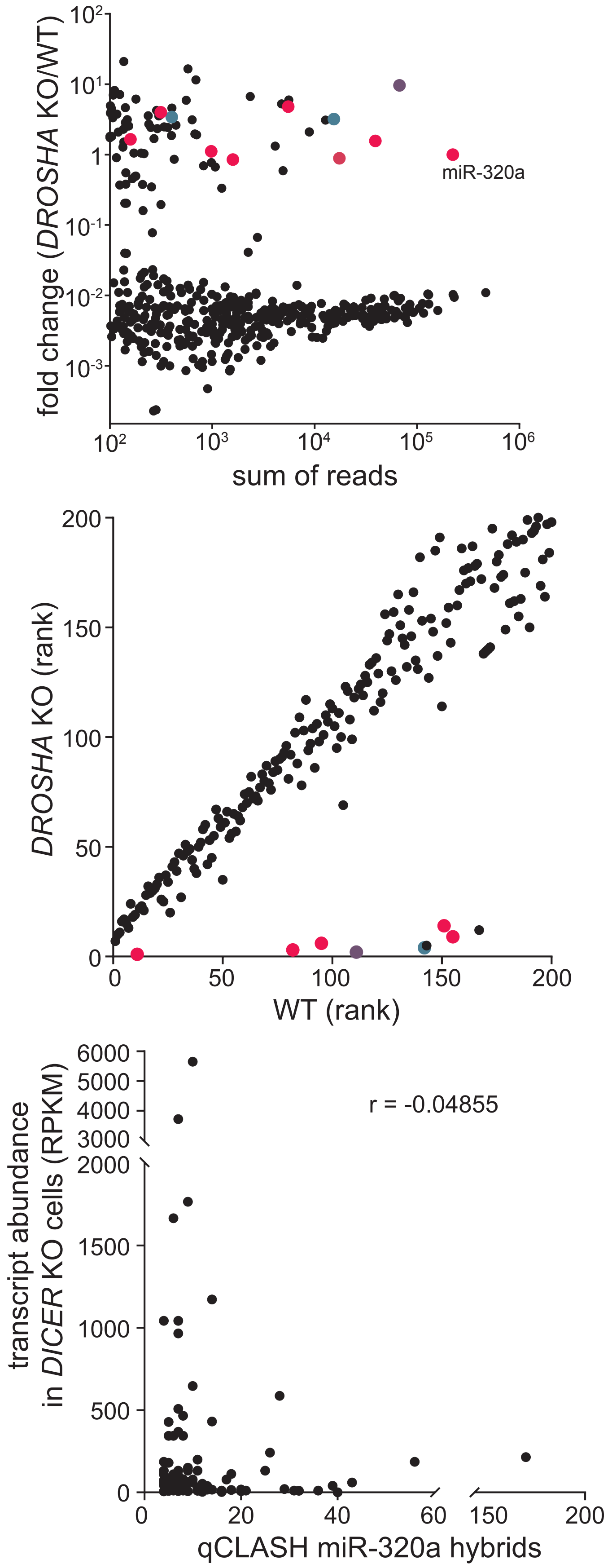

$10^2$
$10^1$
1
$10^{-1}$
$10^{-2}$
$10^{-3}$
fold change (*DROSHA* KO/WT)
miR-320a
$10^2$ $10^3$ $10^4$ $10^5$ $10^6$
sum of reads
200
150
100
50
0
*DROSHA* KO (rank)
0 50 100 150 200
WT (rank)
6000
5000
4000
3000
2000
1500
1000
500
0
transcript abundance in *DICER* KO cells (RPKM)
r = -0.04855
0 20 40 60 150 200
qCLASH miR-320a hybrids

### Figure S4

**A**

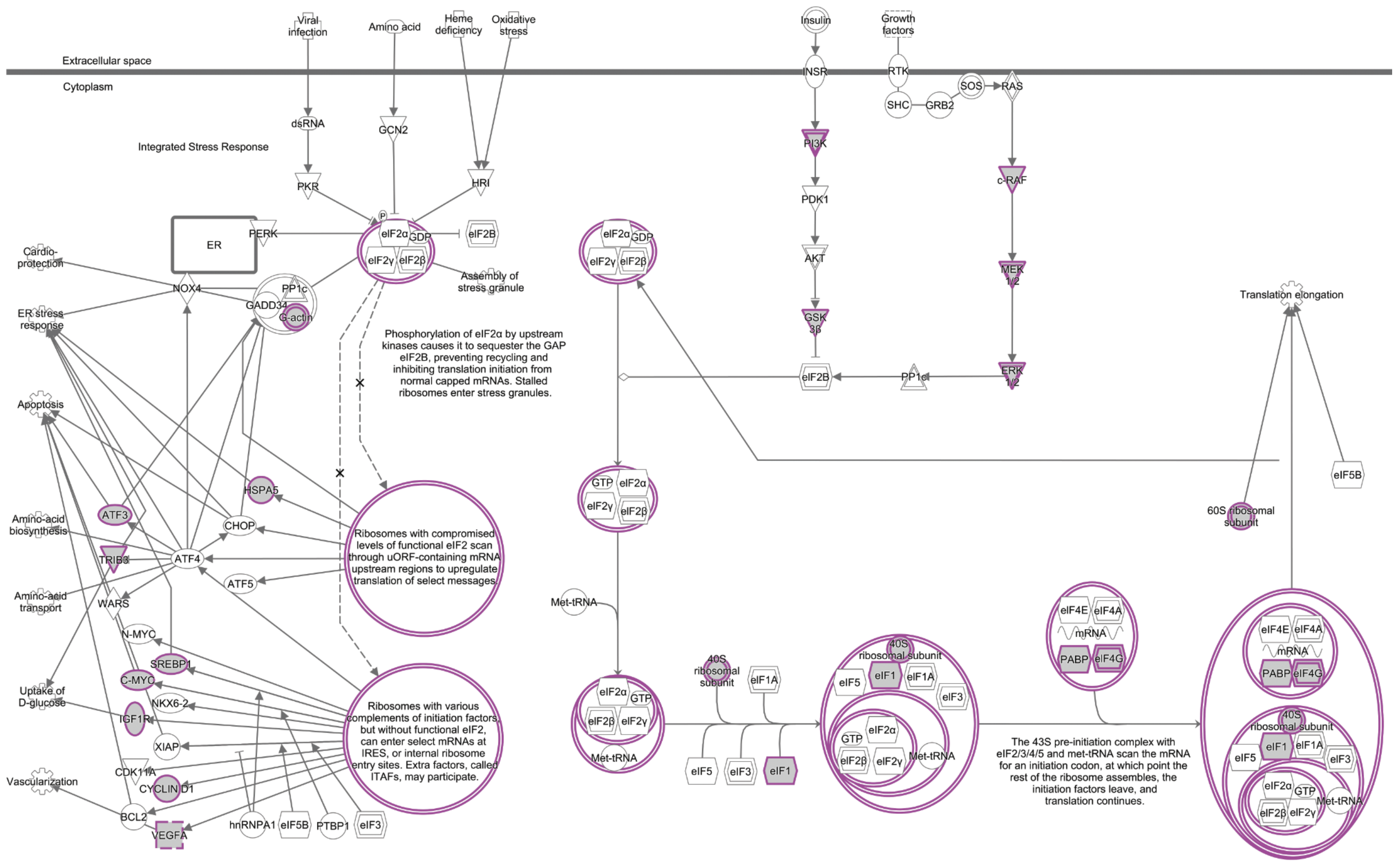

Viral infection
Amino acid
Heme deficiency
Oxidative stress
Insulin
Growth factors
Extracellular space
Cytoplasm
INSR
RTK
SOS
RAS
SHC
GRB2
dsRNA
GCN2
Integrated Stress Response
PI3K
PKR
HRI
c-RAF
PDK1
ER
PERK
eIF2α
GDP
eIF2B
eIF2γ
eIF2β
Cardio-protection
Assembly of stress granule
AKT
MEK 1/2
NOX4
PP1c
GADD34
G-actin
ER stress response
Translation elongation
GSK 3β
Phosphorylation of eIF2α by upstream kinases causes it to sequester the GAP eIF2B, preventing recycling and inhibiting translation initiation from normal capped mRNAs. Stalled ribosomes enter stress granules.
ERK 1/2
PP1c
Apoptosis
GTP
eIF5B
HSPA5
60S ribosomal subunit
ATF3
Amino-acid biosynthesis
CHOP
Ribosomes with compromised levels of functional eIF2 scan through uORF-containing mRNA upstream regions to upregulate translation of select messages
TRIB3
ATF4
ATF5
Amino-acid transport
WARS
Met-tRNA
eIF4E
eIF4A
mRNA
PABP
eIF4G
N-MYC
SREBP1
C-MYC
40S ribosomal subunit
eIF1A
eIF5
eIF1
eIF3
Uptake of D-glucose
NKX6-2
Ribosomes with various complements of initiation factors, but without functional eIF2, can enter select mRNAs at IRES, or internal ribosome entry sites. Extra factors, called ITAFs, may participate.
IGF1R
XIAP
The 43S pre-initiation complex with eIF2/3/4/5 and met-tRNA scan the mRNA for an initiation codon, at which point the rest of the ribosome assembles, the initiation factors leave, and translation continues.
CDK11A
Vascularization
CYCLIN D1
BCL2
VEGFA
hnRNPA1
PTBP1


**B**

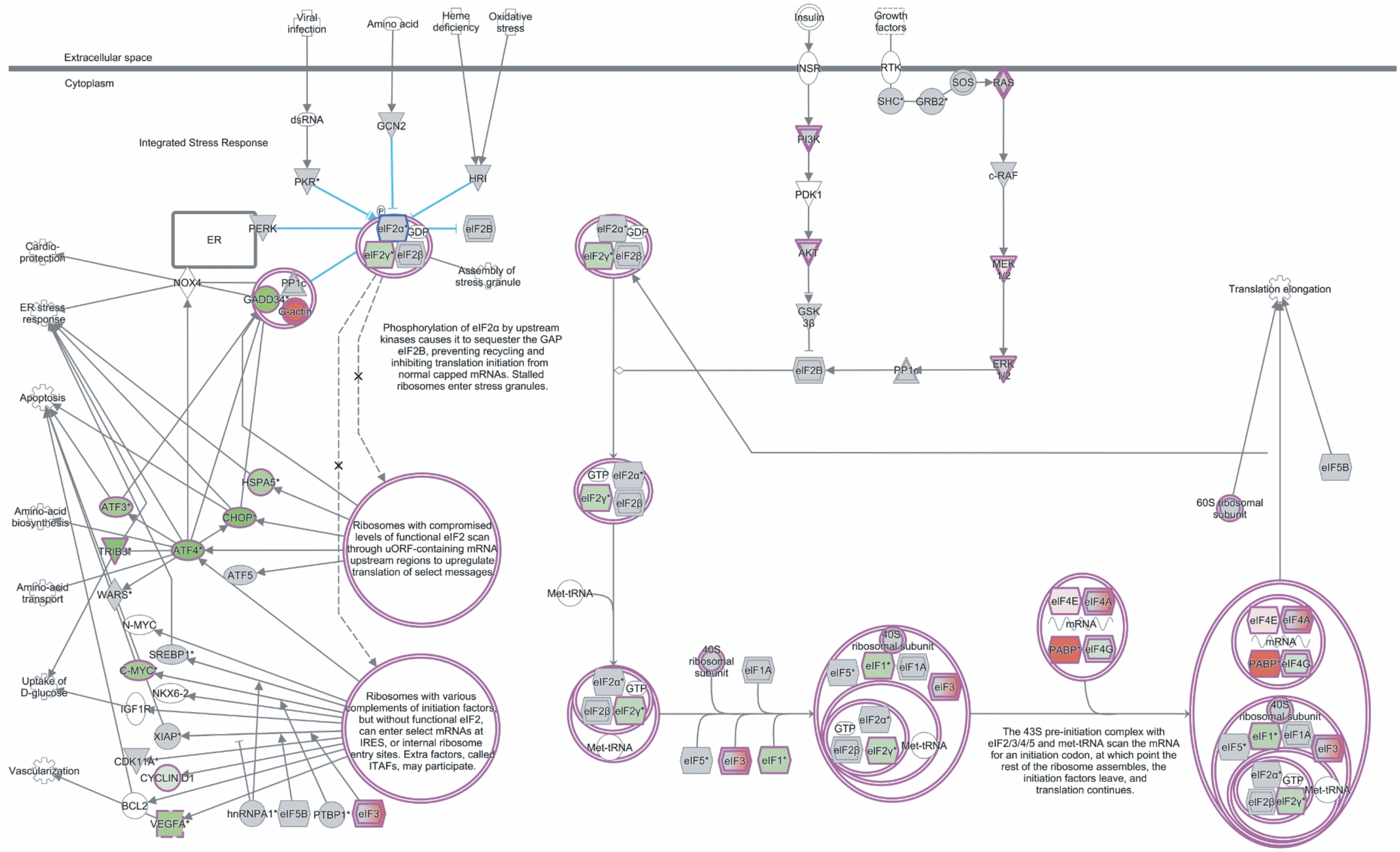

Viral infection
Amino acid
Heme deficiency
Oxidative stress
Insulin
Growth factors
Extracellular space
Cytoplasm
INSR
RTK
SOS
RAS
SHC*
GRB2*
dsRNA
GCN2
Integrated Stress Response
PI3K
PKR*
HRI
c-RAF
PDK1
ER
PERK
eIF2α*
GDP
eIF2B
eIF2γ*
eIF2β
Cardio-protection
Assembly of stress granule
AKT
MEK 1/2
NOX4
PP1c
GADD34*
G-actin
ER stress response
Translation elongation
GSK 3β
Phosphorylation of eIF2α by upstream kinases causes it to sequester the GAP eIF2B, preventing recycling and inhibiting translation initiation from normal capped mRNAs. Stalled ribosomes enter stress granules.
ERK 1/2
PP1c
Apoptosis
GTP
eIF5B
HSPA5*
60S ribosomal subunit
ATF3*
Amino-acid biosynthesis
CHOP*
Ribosomes with compromised levels of functional eIF2 scan through uORF-containing mRNA upstream regions to upregulate translation of select messages
TRIB3*
ATF4*
ATF5
Amino-acid transport
WARS*
Met-tRNA
eIF4E
eIF4A
mRNA
PABP*
eIF4G
N-MYC
SREBP1*
C-MYC*
40S ribosomal subunit
eIF1A
eIF5*
eIF1*
eIF3
Uptake of D-glucose
NKX6-2
Ribosomes with various complements of initiation factors, but without functional eIF2, can enter select mRNAs at IRES, or internal ribosome entry sites. Extra factors, called ITAFs, may participate.
IGF1R
XIAP*
The 43S pre-initiation complex with eIF2/3/4/5 and met-tRNA scan the mRNA for an initiation codon, at which point the rest of the ribosome assembles, the initiation factors leave, and translation continues.
CDK11A*
Vascularization
CYCLIN D1
BCL2
VEGFA*
hnRNPA1*
PTBP1*

### Figure S5

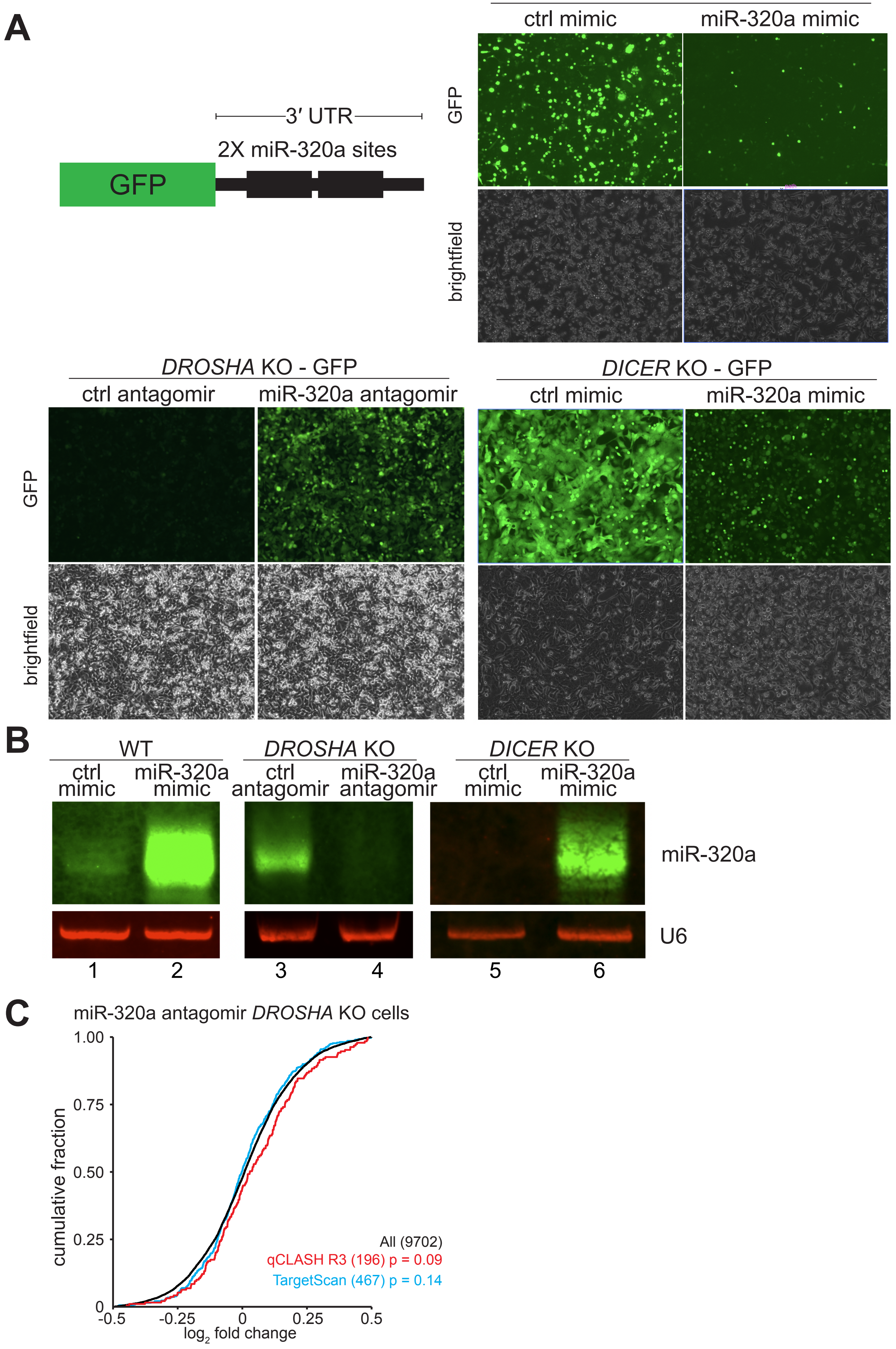
A
WT - GFP
ctrl mimic
miR-320a mimic
GFP
brightfield
3′ UTR
2X miR-320a sites
GFP
DROSHA KO - GFP
ctrl antagomir
miR-320a antagomir
DICER KO - GFP
ctrl mimic
miR-320a mimic
B
WT
DROSHA KO
DICER KO
ctrl mimic
miR-320a mimic
ctrl antagomir
miR-320a antagomir
miR-320a
U6
1
2
3
4
5
6
C
miR-320a antagomir DROSHA KO cells
cumulative fraction
1.00
0.75
0.50
0.25
0
-0.5
-0.25
0
0.25
0.5
log2 fold change
All (9702)
qCLASH R3 (196) p = 0.09
TargetScan (467) p = 0.14

### Figure S6

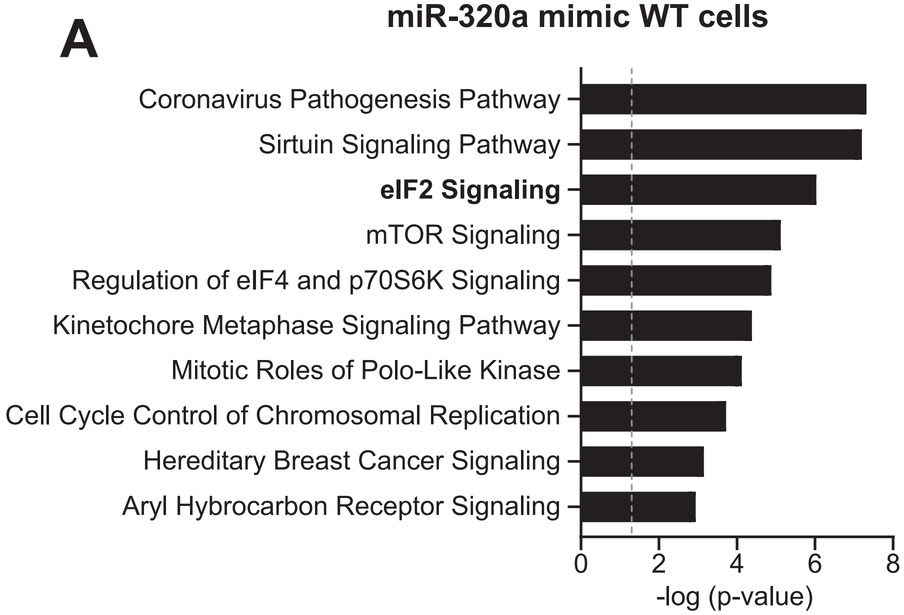
A
miR-320a mimic WT cells
Coronavirus Pathogenesis Pathway
Sirtuin Signaling Pathway
eIF2 Signaling
mTOR Signaling
Regulation of eIF4 and p70S6K Signaling
Kinetochore Metaphase Signaling Pathway
Mitotic Roles of Polo-Like Kinase
Cell Cycle Control of Chromosomal Replication
Hereditary Breast Cancer Signaling
Aryl Hybrocarbon Receptor Signaling
0
2
4
6
8
-log (p-value)

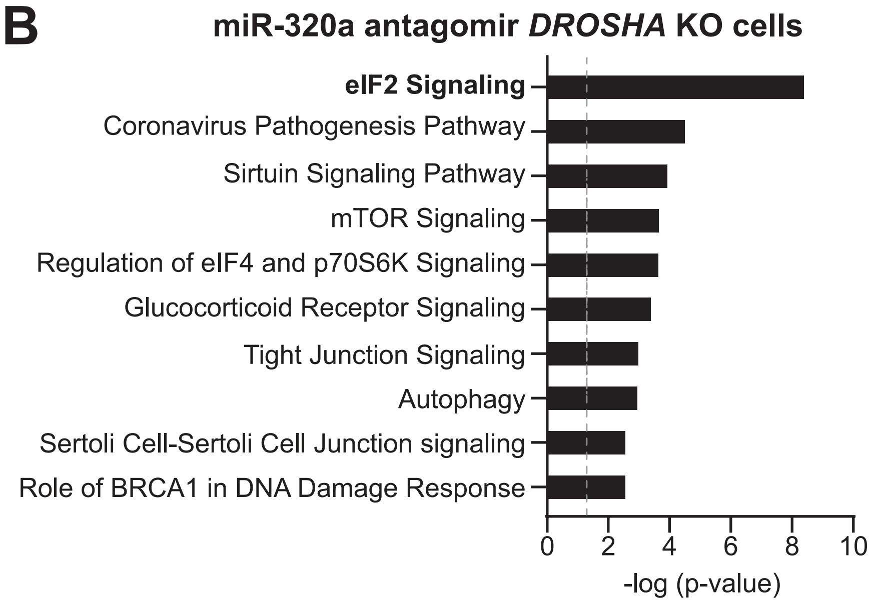
B
miR-320a antagomir DROSHA KO cells
eIF2 Signaling
Coronavirus Pathogenesis Pathway
Sirtuin Signaling Pathway
mTOR Signaling
Regulation of eIF4 and p70S6K Signaling
Glucocorticoid Receptor Signaling
Tight Junction Signaling
Autophagy
Sertoli Cell-Sertoli Cell Junction signaling
Role of BRCA1 in DNA Damage Response
0
2
4
6
8
10
-log (p-value)

### Figure S7

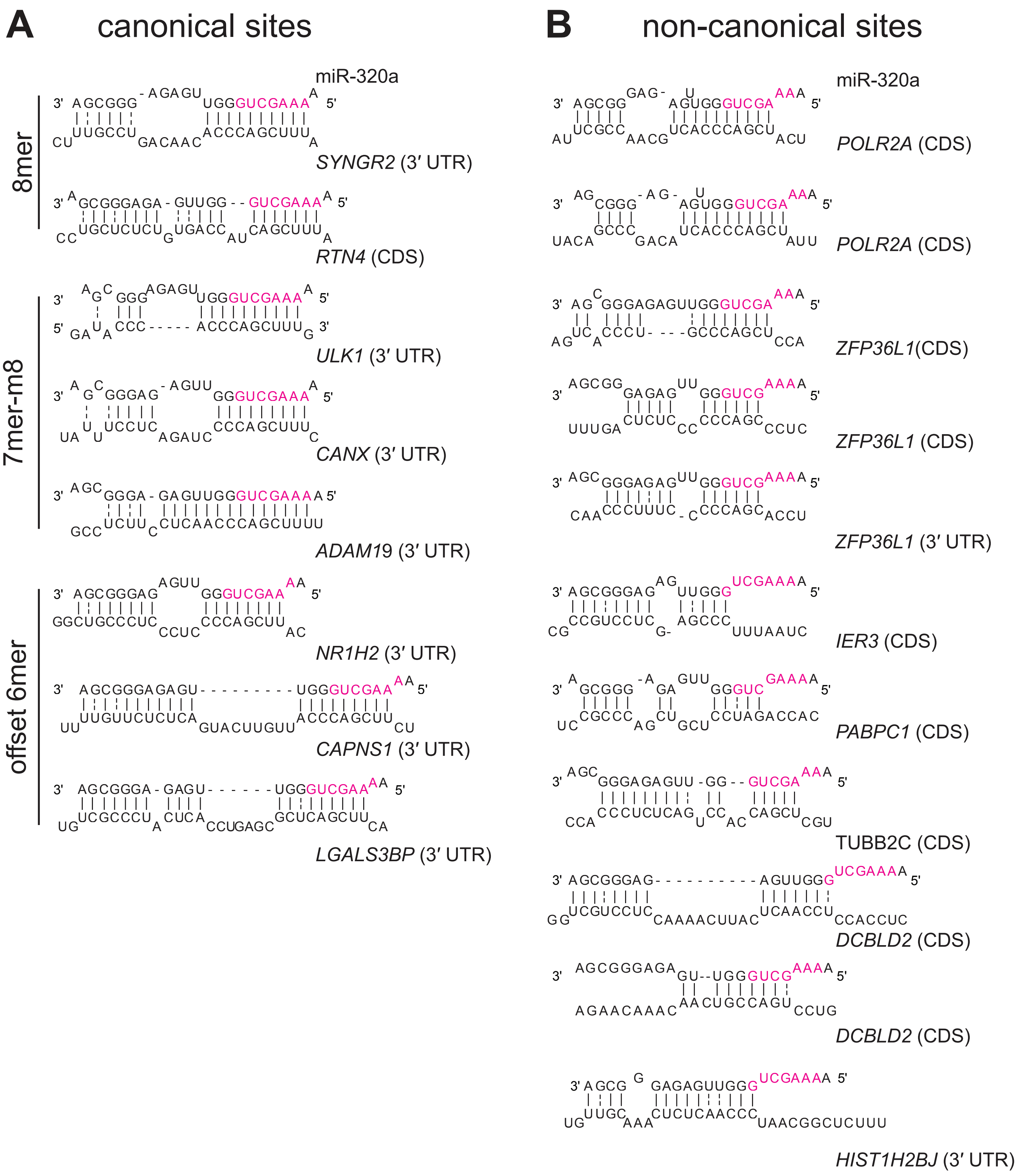
A
canonical sites
B
non-canonical sites
miR-320a
8mer
SYNGR2 (3′ UTR)
RTN4 (CDS)
7mer-m8
ULK1 (3′ UTR)
CANX (3′ UTR)
ADAM19 (3′ UTR)
offset 6mer
NR1H2 (3′ UTR)
CAPNS1 (3′ UTR)
LGALS3BP (3′ UTR)
miR-320a
POLR2A (CDS)
POLR2A (CDS)
ZFP36L1(CDS)
ZFP36L1 (CDS)
ZFP36L1 (3′ UTR)
IER3 (CDS)
PABPC1 (CDS)
TUBB2C (CDS)
DCBLD2 (CDS)
DCBLD2 (CDS)
HIST1H2BJ (3′ UTR)

### Figure S8

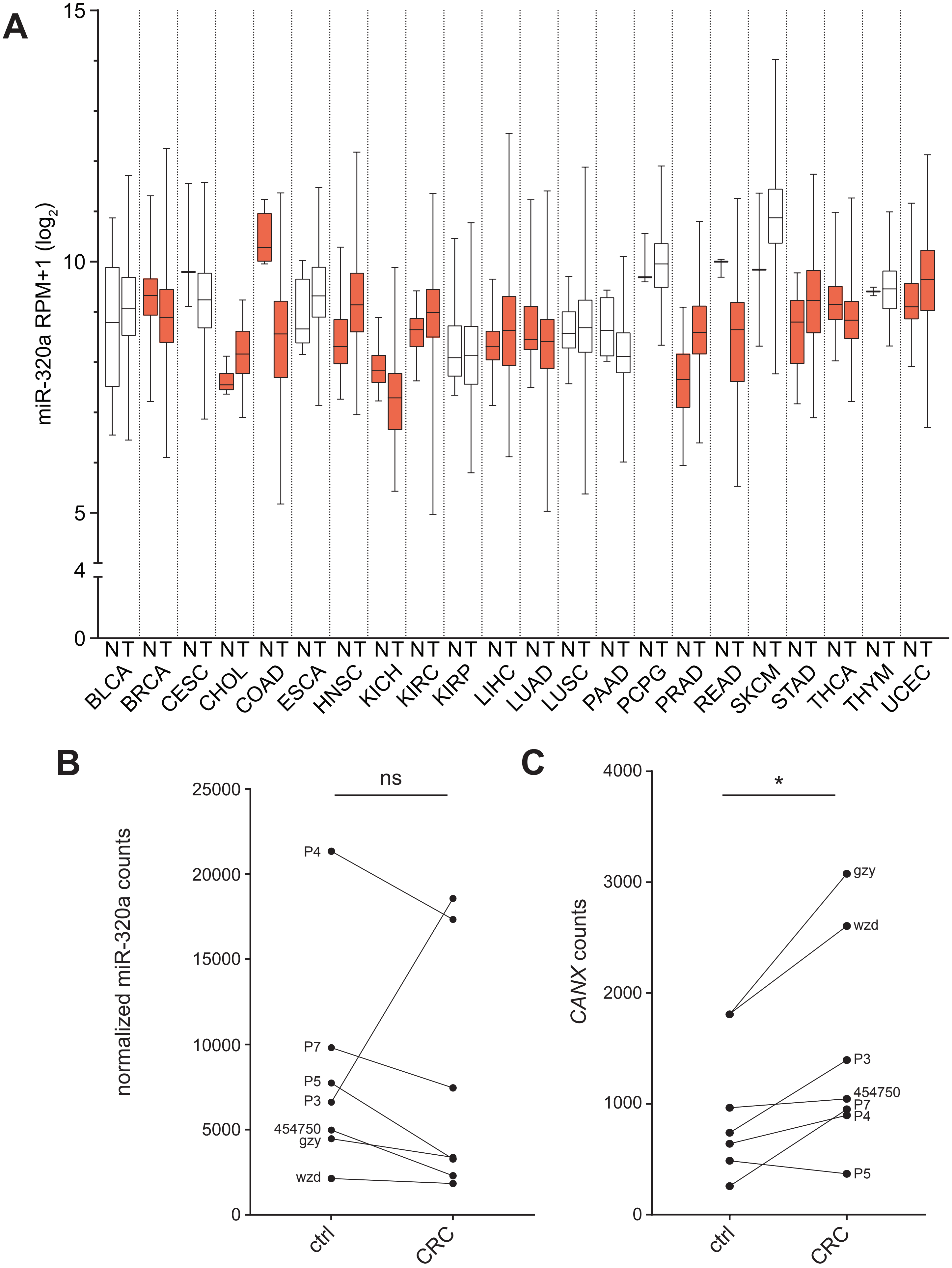

A
miR-320a RPM+1 (log2)
15
10
5
4
0
N T
BLCA
BRCA
CESC
CHOL
COAD
ESCA
HNSC
KICH
KIRC
KIRP
LIHC
LUAD
LUSC
PAAD
PCPG
PRAD
READ
SKCM
STAD
THCA
THYM
UCEC
B
normalized miR-320a counts
25000
20000
15000
10000
5000
0
ns
P4
P7
P5
P3
454750
gzy
wzd
ctrl
CRC
C
CANX counts
4000
3000
2000
1000
0
*
gzy
wzd
P3
454750
P7
P4
P5
ctrl
CRC

### Figure S9

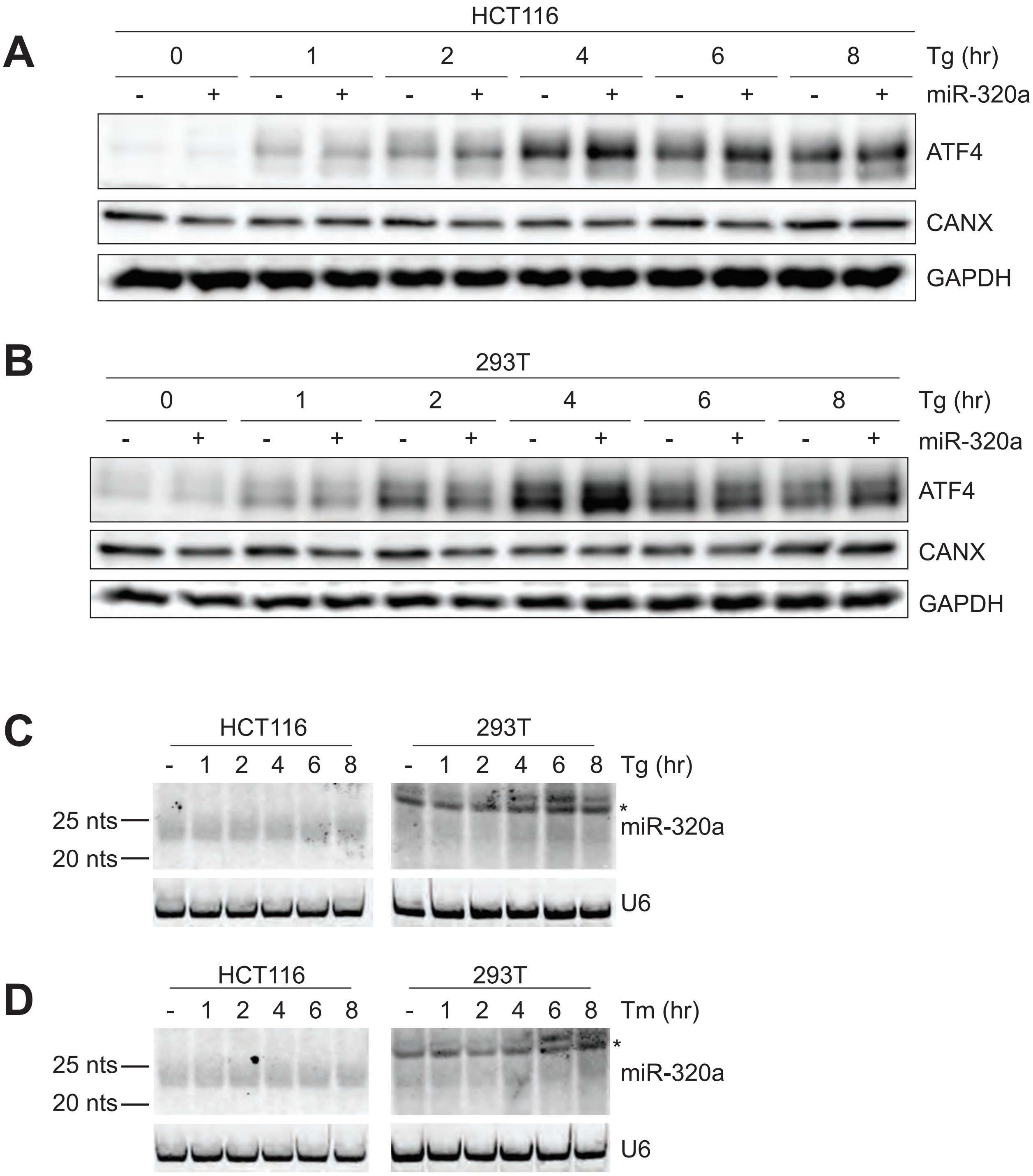

A
HCT116
0 1 2 4 6 8 Tg (hr)
- + - + - + - + - + - + miR-320a
ATF4
CANX
GAPDH
B
293T
0 1 2 4 6 8 Tg (hr)
- + - + - + - + - + - + miR-320a
ATF4
CANX
GAPDH
C
HCT116
- 1 2 4 6 8
293T
- 1 2 4 6 8 Tg (hr)
25 nts
20 nts
*
miR-320a
U6
D
HCT116
- 1 2 4 6 8
293T
- 1 2 4 6 8 Tm (hr)
25 nts
20 nts
*
miR-320a
U6

### Figure S10

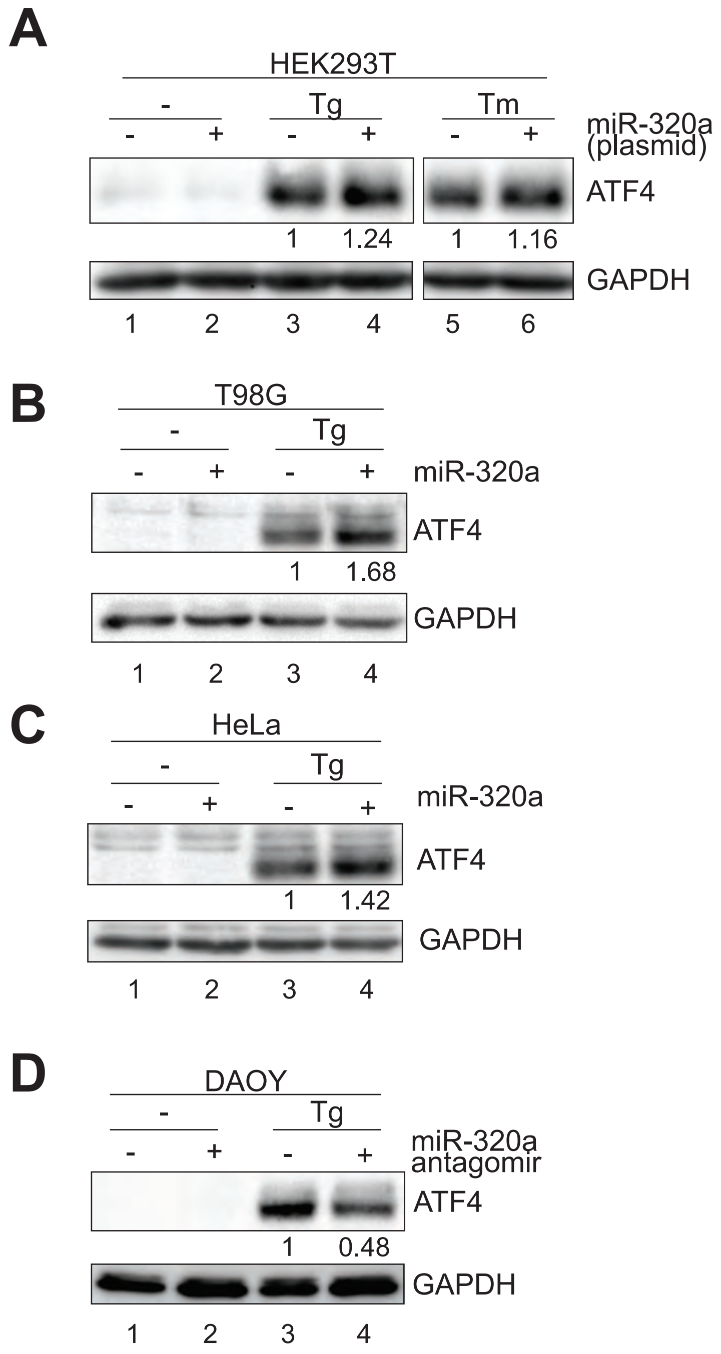
A
HEK293T
-
Tg
Tm
miR-320a
(plasmid)
- + - + - +
ATF4
1 1.24 1 1.16
GAPDH
1 2 3 4 5 6
B
T98G
-
Tg
- + - +
miR-320a
ATF4
1 1.68
GAPDH
1 2 3 4
C
HeLa
-
Tg
- + - +
miR-320a
ATF4
1 1.42
GAPDH
1 2 3 4
D
DAOY
-
Tg
- + - +
miR-320a
antagomir
ATF4
1 0.48
GAPDH
1 2 3 4
